# Selection of streptococcal glucan-binding protein C specific DNA aptamers to inhibit biofilm formation

**DOI:** 10.1101/2024.08.23.608024

**Authors:** Ákos Harkai, Yoon Kee Beck, Anna Tory, Tamás Mészáros

## Abstract

*Streptococcus mutans* is a commensal bacterium in the oral cavity, but its overgrowth leads to biofilm formation and dental caries. *S. mutans* glucan-binding protein C (GbpC) is a cornerstone of the biofilm scaffold and thus a rational target for biofilm inhibition. Here we present a selection of DNA aptamers for GbpC to aid in the development of biofilm inhibiting drugs.

During SELEX, we used the extracellular domain of GbpC as the target of selection and the structurally homologous antigen I/II protein and a GbpC-deficient strain of *S. mutans* as counter-targets. The aptamer candidates obtained were panned using a method based on primer-blocked asymmetric PCR and AlphaScreen. The interaction between the most promising candidates of panning and GbpC was analysed by biolayer interferometry and microscale thermophoresis. Finally, we tested the biofilm inhibitory effect of the aptamers on a wild-type and a GbpC-deficient strain of *S. mutans*. The measurements demonstrated both the efficacy and selectivity of the aptamers. Two of the aptamers studied reduced biofilm formation by 30% in the case of wild-type strains and no reduction was detected in the GbpC-deficient strain, confirming the suitability of the selection strategy presented and providing lead molecules for the development of biofilm inhibitors.

## 1. Introduction

Biofilms are a widespread survival tool for most prokaryotic and some eukaryotic microorganisms. These complex structures consist of diverse extracellular polymeric substances, such as structural nucleic acids, connecting proteins, adhering lipids and scaffolding polysaccharides [1,2]. Using this battery of macromolecules, biofilm-producing cells create a beneficial network and protected milieu for themselves, regulated by a highly sophisticated system, called quorum sensing [3,4]. These heterogeneous communities are more tolerant to the prevailing conditions and often become resistant to antimicrobial treatments [1,5–8]. Therefore, the development of biofilm-inhibiting and biofilm-disrupting molecules are reasonable strategies to reduce the virulence of pathogenic strains.

*Streptococcus mutans* is a gram-positive, facultative anaerobic, predominantly commensal bacterium that is a common and regular biofilm-forming member of the normal microflora in the human oral cavity. However, its overgrowth contributes to the formation of dental plaque on the tooth surface and, in turn, to the onset of oral diseases, such as dental caries. Among its virulence factors, glucan-binding proteins (Gbps) are key players in biofilm architecture and are essential for establishing connections between cells and scaffolding polysaccharides [9–15]. The cell wall-anchored GbpC of *S. mutans* has received particular attention, as its X-ray crystal structure and molecular interactions have been thoroughly described [16]. The authors compared the properties of GbpC with those of streptococcal antigen I/II (AgI/II), which shows structural similarity but does not bind dextran [16]. At both morphological and molecular level, the function of GbpC has been described that is crucial for the extensive formation of sucrose-dependent biofilms [16–18]. In light of these findings, GbpC appears to be a promising target for inhibition biofilm formation in *S. mutans* strains.

Although recent reports suggest that the presence of a surface is not essential for the initiation of biofilm formation [19], most disease-causing populations still colonise on the surfaces of tissues, wounds, implants, medical devices or other everyday tools [20–23]. To overcome this issue, a variety of molecules have been identified, that can influence either the production or the structure of the maturing biofilm. For example, some sweeteners can alter the expression of biofilm-related genes in certain prokaryotes, thereby reducing biofilm formation [24,25]. Antimicrobial peptides have also been shown to disrupt the cell membrane of biofilm-forming bacteria [26,27]. In addition, numerous antibodies have been developed to combat biofilm-associated infections [28–32]. Despite the intensive research, no standard anti-biofilm treatment has yet been developed.

Aptamers are artificial RNA or DNA oligonucleotides that can theoretically bind specifically to any type of target molecule, from small molecules to cells. They have undergone dynamic development since their first publication [33,34], with numerous examples demonstrating both their diagnostic and therapeutic potential [35–41]. Despite the favourable properties of aptamers [42], their routine use in diagnostics and therapeutics is still awaited. Recently, only the second aptamer, *avacincaptad pegol*, which binds to and inhibits the complement protein C5 has been approved for the treatment of advanced form of age-related macular degeneration [41,43]. According to the scientific literature, only about a dozen bacteriostatic aptamers have been generated since the first aptamer was described [4,44], indicating that the antibacterial potential of aptamers has not yet been fully exploited. Most bacteriostatic and bacterial diagnostic aptamers have been selected by cell-SELEX using bacterial cells [4,44]. Biofilm inhibitory aptamers generated against whole cells include *Proteus mirabilis*, an enteropathogenic *E. coli* stain (EPEC K1.1) and *S. mutans* selective aptamers [45–47]. The protein targeting SELEXs were primarily designed to reduce biofilm formation by disrupting the quorum sensing system of the pathogens, using the proteins involved as targets for selection, such as Salmonella Invasion Protein A (SipA) and the C4-HSL signalling molecule of *Pseudomonas aeruginosa* [48,49].

In this study, we present the selection of the first GbpC-specific aptamers with biofilm inhibitory potential to date. To this end, we produced the target of selection, the extracellular domain of GbpC, by bacterial overexpression and demonstrated its correct folding by various methods. In our selection protocol, the target protein was immobilised on magnetic beads and counter-selection steps were introduced using AgI/II protein and GbpC-deficient (ΔGbpC) *S. mutans* cells. The aptamers obtained were analysed by different approaches to demonstrate their specificity and to determine their affinity for GbpC. Finally, we showed that the best performing aptamers reduced biofilm formation by *Streptococcus mutans* UA130 strain.

## 2. Materials and methods

### 2.1. Molecular cloning and bacterial strains

The gene fragments for streptococcal cell surface protein expression were designed based on a study of Mieher et al. [16], chemically synthesised and provided in pUCIDT-Amp vector constructs (IDT). The coding sequences were transferred into the N-terminal 6xHis-tagging pET-28a expression vector (Novagen) using NdeI and XhoI restriction endonucleases and T4 DNA Ligase (Thermo). The assembled constructs containing either GbpC^(111-522)^ or AgI/II^(457-993)^ gene fragments were purified and sequenced (Macrogen).

*Streptococcus mutans* UA130 (wild-type) and GbpC deficient mutant (ΔGbpC) strains [18] were cultured either on BHI agar plates or in BHI media (Jena Bioscience) and were grown at 37°C in an aerobic incubator.

### 2.2. Overexpression and purification of recombinant proteins

To overexpress GbpC (50 kDa) and AgI/II (65 kDa) protein fragments, an auto-inducing protein production approach was used [50]. First, the appropriate vector holding *E. coli* BL21 (DE3) colonies were added to 2 ml MDG non-inducing medium supplemented with 50 µg/ml kanamycin. The cell cultures were grown at 37°C for 5 hours with 220 rpm rotation in an aerobic incubator. Then, 50 µl from each non-inducing culture was inoculated into 100 ml ZYM-5052 auto-inducing medium supplemented with 50 µg/ml kanamycin. The induced broths were grown aerobically at 37°C for 16 hours with 220 rpm rotation. Afterwards, 100 ml of auto-induced mediums were split into 5 sterile conical tubes and centrifuged at speed of 4000 rpm for 20 min at 4°C. The resulting bacterial pellets were stored in a freezer until the purification process.

Recombinant 6xHis-tagged proteins were purified using PureCube INDIGO Ni-MagBeads (Cube Biotech) according to manufacturer’s protocol. For SELEX, the purified proteins were kept on paramagnetic beads and eluted, then buffer exchanged to 1x Artificial Saliva Buffer (ASB) using 30K Amicon Ultra filter (Merck) for binding assays (components of 1xASB are listed in Table S1). The purity of the protein fractions was verified by SDS-PAGE.

### 2.3. SELEX procedure

The single-stranded DNA aptamer library was obtained from Sigma. The initial library consists of a 40 nucleotide comprises variable region that is flanked by primer regions: 5’-AGATACCAATACGCTGCC-(N40)-GCCACTGGTAACGACATC-3’ (approx. 23.5 kDa). Prior to the selection cycles, the aptamer library was denatured in nuclease-free distilled water at 95°C for 5 min and then cooled to room temperature. The selection procedure was a combination of the FluMag-SELEX [51,52], which used the magnetic bead-immobilised GbpC protein for positive selection steps and AgI/II for negative selection steps, moreover the cell-SELEX [53], which used ΔGbpC *S. mutans* cells. The selection pressure was gradually increased to eliminate the non-specific and weakly binding oligonucleotides (see detailed SELEX protocol in Table S2). At the end of the positive selection rounds, the oligonucleotides bound to the target molecule were eluted with nuclease-free distilled water using a thermoblock at 95°C for 5 min. This elution served as a template for the following PCR step.

Initially, 1 nmol of aptamer library was added to 600 pmol paramagnetic bead coupled GbpC target molecule and these were incubated in 1.3 ml 1xASB (see components in Table S1) at room temperature using orbital rotator. As competitors, mucin (Pickering) and salmon sperm DNA (Sigma) were added, and incubations were followed by various washing steps as detailed in Table S2.

To minimize by-product formation, emulsion PCR was conducted to amplify the enriched oligonucleotide pool of selection cycles. The water-in-oil reaction mixture was compiled according to manufacturer’s protocol (EURx, Micellula Kit). 50 µl water-fraction contained 1x buffer for KOD XL DNA polymerase, 0.02 U/µl KOD XL DNA polymerase (Toyobo), 0.2 mM (each) KOD dNTPs, 0.3-0.3 µM unlabelled forward and 5’-biotinylated reverse primers, 4% DMSO, 0.02 µg/µl BSA and template DNA. PCR conditions were as follows: an initial denaturation at 94°C for 5 min was followed by 25 thermocycles of 94°C for 30 sec, 50°C for 5 sec, 72°C for 20 sec and final extension of 72°C for 5 min. Emulsion PCR products were purified using a scaled-up protocol of OligoClean and Concentrator Kit (Zymo Research). The resulting PCR products were analysed by 10% polyacrylamide gel electrophoresis after staining with GelGreen® dye (Biotium) (all oligonucleotide used are listed in the Table S3).

The cleaned double-stranded ePCR products were coupled to streptavidin-coated paramagnetic beads following manufacturer’s protocol (Dynabeads M-270 Streptavidin, Thermo). The positive strand was separated by alkalic denaturation using 50 µl of freshly prepared 50 mM NaOH for 10 minutes. The denatured single strands containing supernatant was neutralized with 7.5 µl of 400 mM NaH_2_PO_4_ and used in the next selection round.

During the ePCR of the last selection cycle, unlabelled primers were added to the reaction mixture and the purified dsDNA products were cloned into TOPO vector using the Zero Blunt TOPO Cloning Kit (Invitrogen). The construct was transformed into α-Select Gold chemically competent cells (Bioline). The insertion of aptamer candidates was confirmed by colony PCR using vector specific primers (Table S3). 55 of the PCR products with the correct size were analysed by Sanger sequencing and the DNA motifs were subsequently evaluated using the MEME Suite online tool [54].

### 2.4. Panning of aptamer candidates

The individual oligonucleotides were screened using the AlphaScreen (PerkinElmer) technique to pinpoint the promising aptamer candidates. To this end, an optimized primer-blocked asymmetric (PBA) PCR [55,56] was applied to produce the appropriate 5’-biotinylated aptamer candidates. 20 µl of PBA-PCR mixture comprised of 1x buffer for KOD XL DNA polymerase, 0.02 U/µl KOD XL DNA polymerase (Toyobo), 0.2 mM (each) KOD dNTPs, 500 nM 5’-biotinylated forward, 25 nM unlabelled reverse and 475 nM 3’-phosphorylated reverse primers, 4% DMSO, and 0.5 µl of 50x diluted colony PCR product as template DNA. The PCR conditions were as follows: an initial denaturation at 94°C for 3 min was followed by 45 thermocycles of 94°C for 30 sec, 50°C for 5 sec, 72°C for 20 sec and final extension of 72°C for 5 min. Following PCR, the complement oligonucleotide of the reverse primer was added to the mixture in a five-fold molar excess of blocked reverse primer (Table S3). The mixture was incubated at 95°C for 5 min then cooled to room temperature to obtain fully single-stranded aptamer candidates [56].

The Histidine Detection Kit (PerkinElmer) was used for ALPHA measurement. The assay was set up in an AlphaPlate 384-SW (PerkinElmer) with a final volume of 18 µl. The reaction mixtures contained either 500, 750, 1000 nM 6xHis-GbpC or 750 nM 6xHis-AgI/II, approximately 12.5 nM of 5’-biotinylated ssDNA produced by PBA-PCR, 20 µg/ml of donor and acceptor beads in 1xASB supplemented with 0.1 mg/ml mucin and 0.1 µg/ml salmon sperm DNA. The protein and the aptamer candidates were pre-incubated for 60 min at 25°C.

Then the nickel-chelate acceptor beads were added, followed by the streptavidin donor beads and incubated for 30-30 min in dark. Chemiluminescent signal was detected by EnSpire multimode plate reader (PerkinElmer).

### 2.5. In vitro binding assays

#### 2.5.1. BLI

Five aptamer candidates from screening were synthesized with 5’-biotin-TEG labelling (Eurofins) for biolayer interferometry (BLI) measurements (Table S3). Working solutions of either aptamer candidates or biotin-dextran (Thermo) at a 1 µM concentration were prepared in 1xASB, then loaded onto separate Super Streptavidin biosensors (Fortebio). The immobilised molecules were then exposed to increasing concentrations (0 -50 µM) of GbpC protein. Loading was followed by measuring the initial baseline (30 sec), the association (90 sec), the dissociation (90 sec), the washing (90 sec) and the final baseline (30 sec) steps. The binding kinetics were examined using the BLItz instrument (Fortebio) and evaluated with BLItz Pro (v1.2) software.

#### 2.5.2. MST

Flanked and truncated forms of two aptamer candidates were synthesized with 5’-Cy5 fluorescent labelling (Eurofins) for microscale thermophoresis (MST) measurements (Table S3). In direct binding assays, the 1xASB-exchanged 200 µM GbpC stock solution was diluted to 3 nM in a 16-step serial dilution, 1:1 (v/v) with 1xASB. Each dilution step was added with an equal volume of either 100 nM Cy5-labelled aptamer candidate or 100 nM FITC-labelled dextran (Sigma) in 1xASB + 0.02% Tween-20 solution. After 5 min incubation at 25°C, reaction mixtures were loaded into Monolith Capillaries (NanoTemper). The LED power was set to 20% for both excitation (red for Cy5, blue for FITC), while the MST power was set to 40%. The fluorescence signal was monitored before, during and after the infrared laser was turned on for 3, 20 and 1 sec, respectively using the Monolith NT.115 (NanoTemper) instrument. The MST traces were then analysed, and dissociation constants (K_D_) were determined with MO.Affinity Analysis (v2.3) software (NanoTemper).

In competitive binding assays, target GbpC was diluted to 4 µM in 1xASB + 0.02% Tween-20 and it was pre-incubated with tracer molecule, with either 100 nM flanked Cy5-A39 or 100 nM flanked Cy5-A96 for 5 min at 25°C. Thereafter, a 16-step serial dilution was prepared in 1xASB using 100 µM FITC-dextran as the ligand. The pre-incubated mixture was then added to an equal volume of each of the dilution series. The red LED power and MST power were set to 20% and 40%, respectively. The fluorescence signal was monitored and analysed as described above, and ‘half maximal effective concentrations’ (EC_50_) of dextran were determined with MO.Affinity Analysis (v2.3) software (NanoTemper).

### 2.6. Biofilm inhibition assay with aptamers

*S. mutans* UA130 wild-type and ΔGbpC strains were cultured in BHI broth until mid-log phase, then they were inoculated into freshly prepared BHI + 1% sucrose biofilm-forming medium (BFM). Next, 98 µl of BFM were dispensed into sterile U-shaped 96-well microtest plate (Sarstedt), and 2 µl of 100 µM unlabelled flanked aptamer (Eurofins) stock solution was added to each well. The microtest plate was incubated for 16 hours at 37°C in an aerobic, static incubator. Aptamer-free (untreated) inoculated BFM samples were used as controls for normalization.

The following day, planktonic cells containing supernatants were discarded and each well was washed three-times with autoclaved 1xPBS. The remaining biofilm structures were stained with 0.05% crystal violet solution and re-suspended in 30% acetic acid. The samples were then transferred into F-shaped 96-well microtest plate (Sarstedt), and the absorbance was measured at 595 nm using CLARIOstar (BMG Labtech) microplate reader instrument. The degree of inhibition effect was calculated according to the equation below.

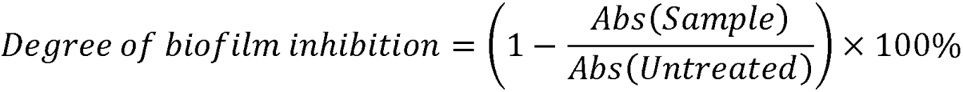

## 3. Results and discussion

### 3.1. Production of recombinant proteins

The cell wall-anchored GbpC of *S. mutans* is essential for glucan-dependent adhesion between the bacterial cells and carbohydrate components of the biofilm [16–18]. Given its key role in the initial stages of biofilm formation and its cell surface-anchored localisation, GbpC represents an optimal target for the selection of aptamers with biofilm inhibitory potential.

To produce the GbpC and AgI/II protein fragments, we used the convenient and highly productive *E. coli* auto-induction system and purified the proteins by immobilised metal affinity chromatography (IMAC) using the hexahistidine tag of the recombinant protein fragment. The production of membrane protein domains can be a challenging task, but we encountered no difficulties with our approach and were able to purify a sufficient amount of soluble proteins (Figure 1A).

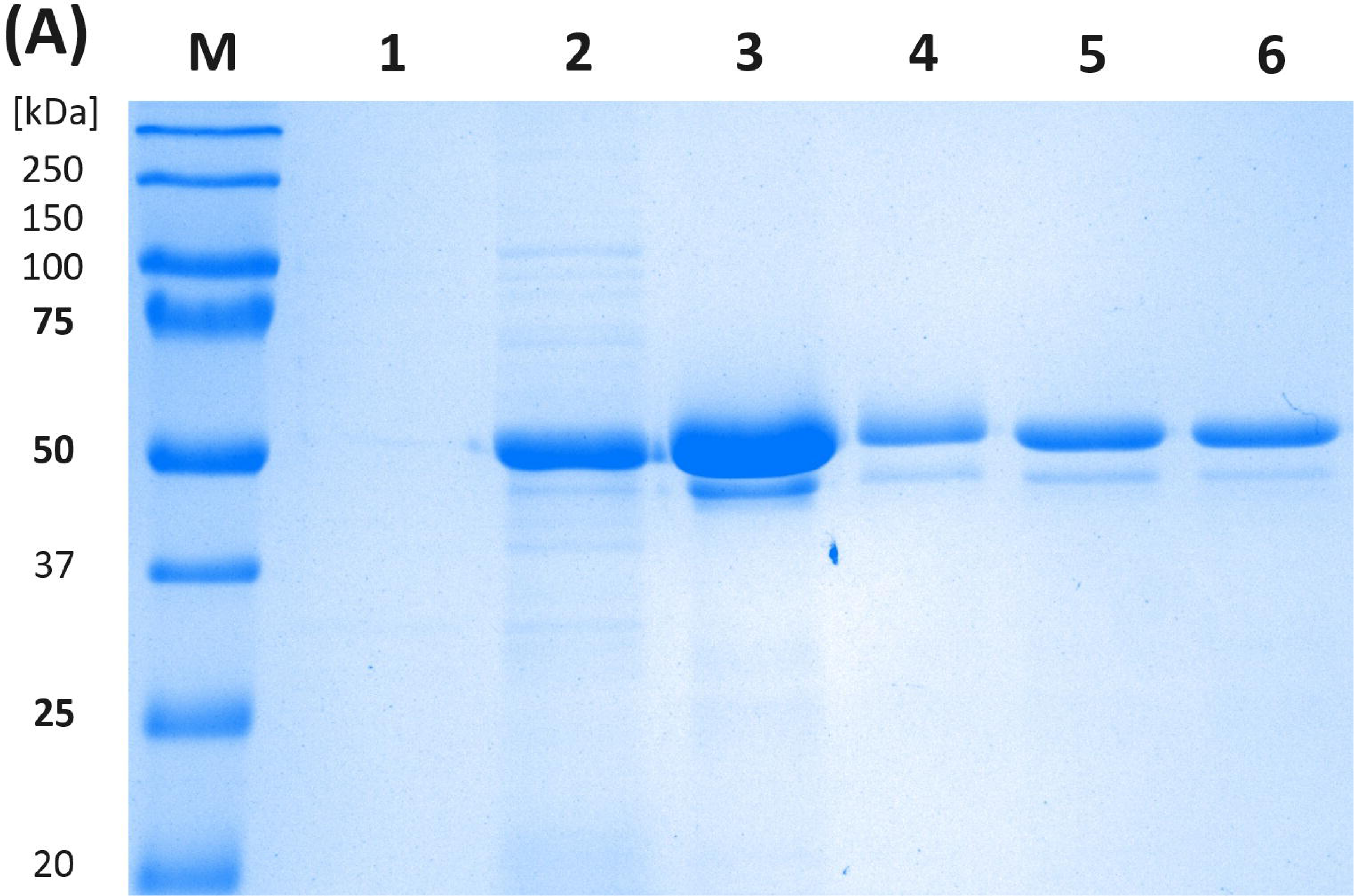
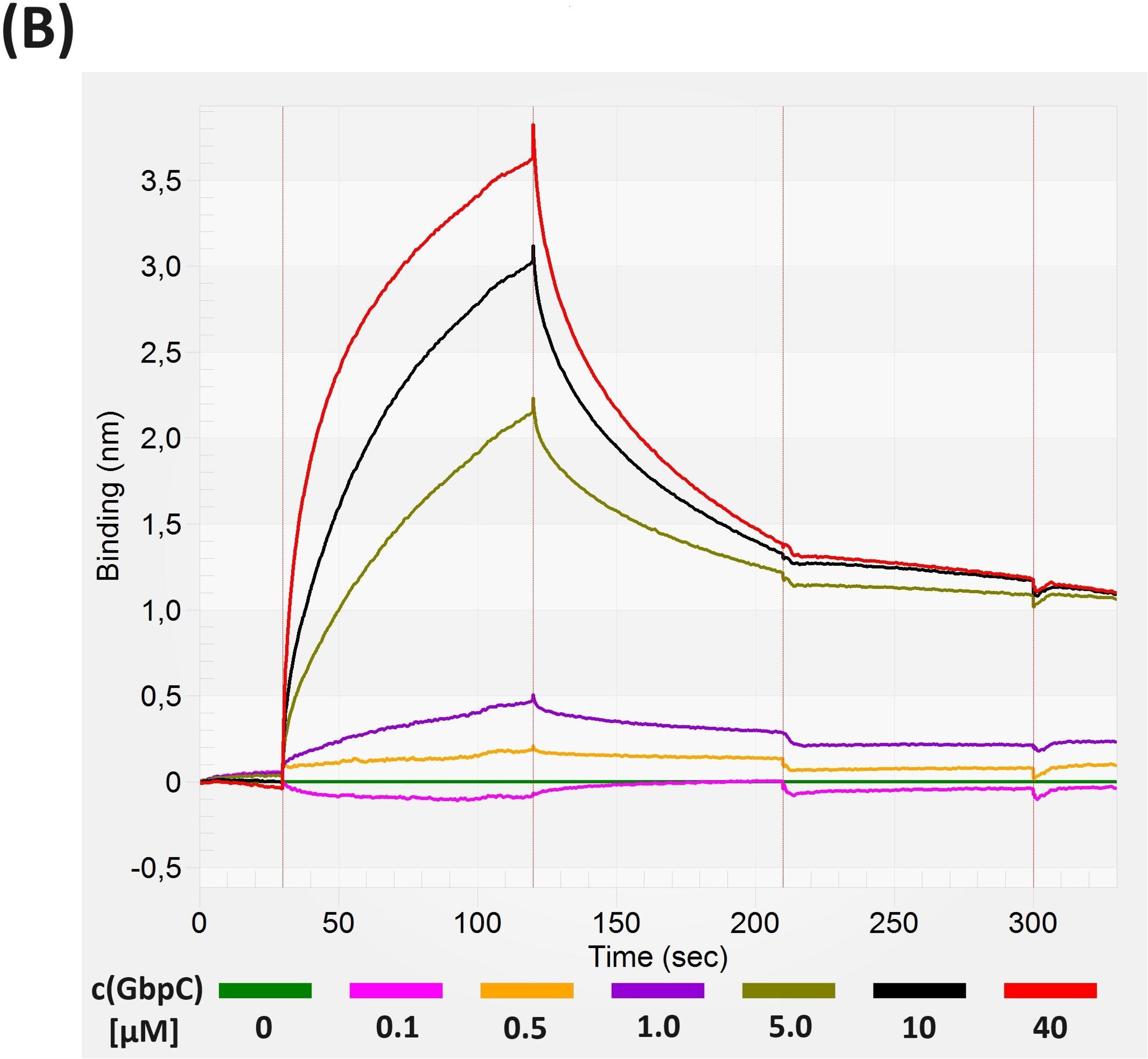
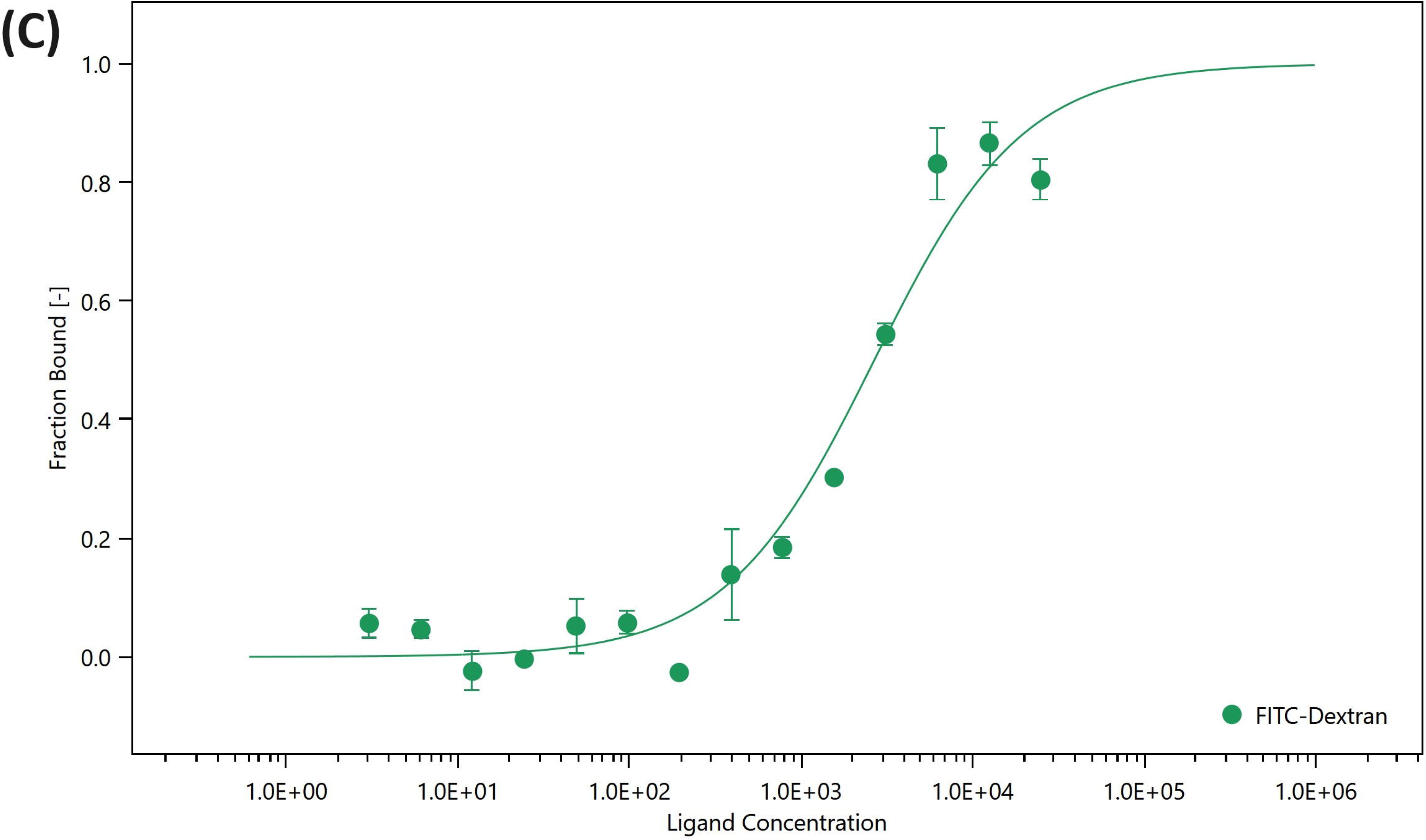

Although the solubility of GbpC suggested that the protein was correctly folded, we also sought to provide experimental evidence that the protein indeed had a native-like spatial structure. To this end, three types of measurements were conducted to ascertain whether the purified protein fragment exhibited the previously published functional properties of GbpC. These characteristics included increased structural stability in the presence of calcium ions and the ability to bind glucan [16]. To demonstrate the enhanced stability of the calcium-bound protein, a thermal unfolding assay was performed using the purified protein in PBS, PBS supplemented with calcium, and a calcium-containing artificial saliva buffer (see Method S1). The presence of calcium in the buffer resulted in the appearance of a distinct additional peak in the melting curve, indicating increased spatial structural stability of GbpC (Figure S1). We applied two different approaches to demonstrate the glucan-binding capacity of the purified protein: biolayer interferometry (BLI) and microscale thermophoresis (MST). The results of both methods were in alignment with previous publications [16], showing a low micromolar dissociation constant for the protein-dextran complexes, which was 13.69 and 2.46 µM when measured by BLI and MST, respectively (Figure 1B and 1C).

Collectively, these data confirmed the native folding of purified GbpC and indicated that it could be used as a target protein for SELEX.

### 3.2. Selection and panning of aptamer candidates

We designed the SELEX conditions taking into account the intended use of the aptamers; therefore, a mucin-enriched artificial saliva was used as a selection buffer (Table S1). To increase the specificity of the selected aptamers, two types of counter-selection were performed during SELEX. On the one hand, we used the 457-993 protein fragment of AgI/II, which shows structural similarity to the 111-522 fragment of GbpC (Figure S2), and on the other hand, a ΔGbpC *S. mutans* strain to eliminate oligonucleotides that bind to bacterial cells via cell wall components other than GbpC (Figure 2). A decreasing amount of magnetic beads with GbpC was used in the successive steps of SELEX and counter-selection steps were introduced after cycles 3, 7 and 5 using AgI/II-coated IMAC magnetic beads and the ΔGbpC mutant *S. mutans* strain, respectively. Further details of selection are given in Table S2.

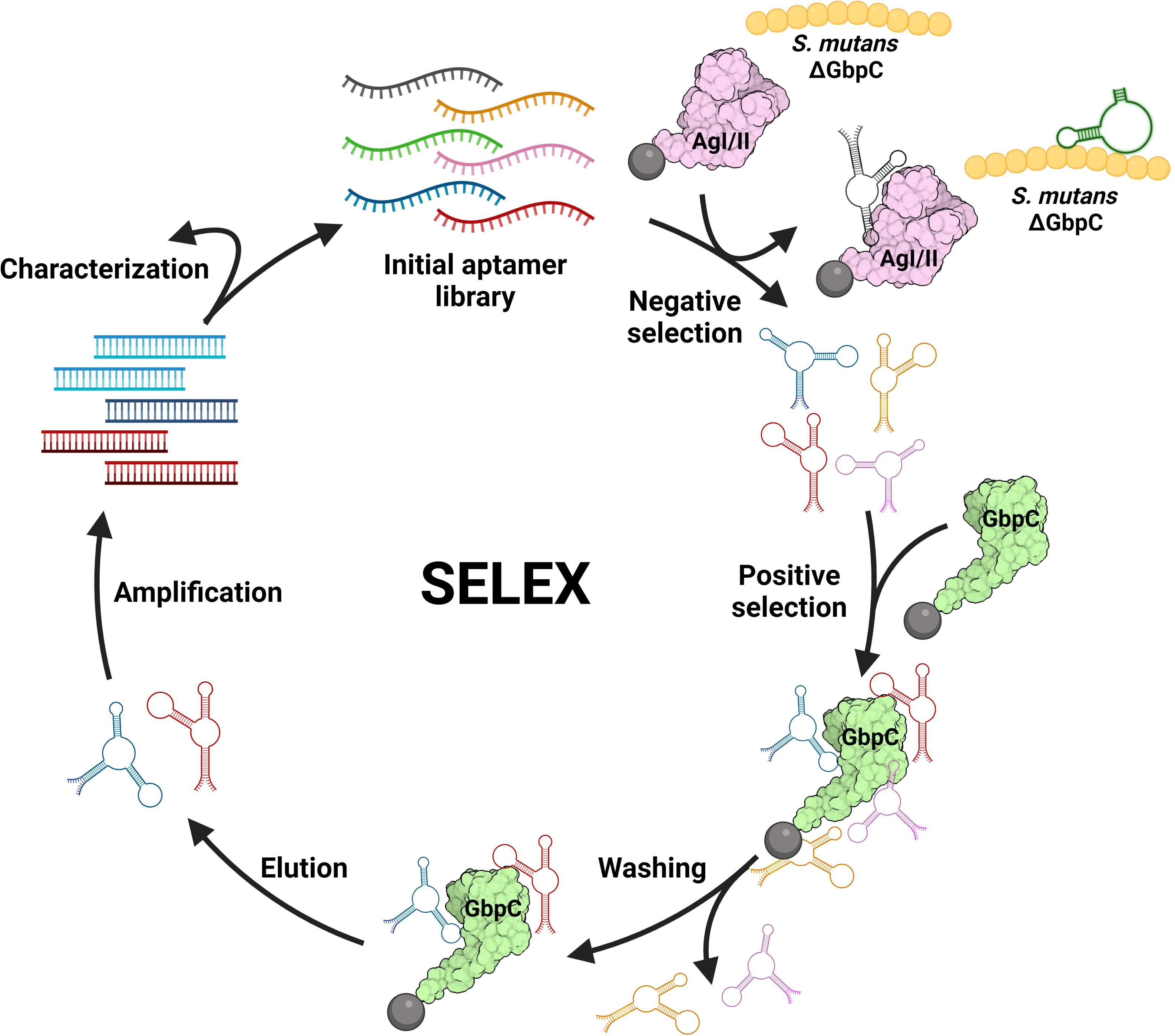

Following the final SELEX cycle, the enriched aptamer library was cloned into the TOPO vector and the nucleic acid sequence of plasmids from 55 bacterial colonies of transformed cells was determined by Sanger sequencing. The results of sequencing proved enrichment of the library. We identified 42 unique sequences among the 55 sequenced aptamer candidates, of which one was presented in 6 copies (A10) and 6 others were identified in 2 copies (A32, A41, A49, A55, A57 and A75). The in-silico analysis also revealed that variable regions containing cytosine bases were highly overrepresented in the oligonucleotides obtained after the final step of the SELEX (see illustration of obtained sequences on Figure S3).

These data demonstrated enrichment of the aptamer library, but did not provide any real information on their target binding properties. Therefore, in order to pinpoint the aptamers most capable of binding to GbpC, we set out to experimentally identify the best candidates in vitro.

Previously, we demonstrated that the Amplified Luminescent Proximity Homogenous Assay (ALPHA) is a simple, cost-effective method for screening aptamer–target protein interactions [57]. We combined this approach with primer-blocked asymmetric PCR (PBA-PCR) to circumvent the complexity of commonly used ssDNA generation protocols and set up the AlphaScreen assay [55,56]. Hexahistidine-tagged GbpC was mixed with biotinylated aptamer candidates produced PBA-PCR and completed by the addition of the AlphaScreen acceptor and donor beads. Measurement of the chemiluminescence signal (Figure 3) showed that about a dozen of the oligonucleotides analysed bound to the target protein of the selection (A29, A30, A38, A39, A40, A41, A47, A65, A74, A94, A96). Notably, GbpC itself bound moderately to the bare donor bead, resulting in a relatively high background signal (dashed lines on Figure 3). This phenomenon can probably be explained by unreleased, patented components of the donor bead (e.g. agarose, cellulose, dextran [58]), to which the well-folded overexpressed GbpC readily binds due to its glucan-binding function (Figure S4). To see if the best performing aptamer candidates shared common nucleic acid sequence motifs, we applied the MEME Suite online tool, which identified only one loose C-enriched motif in 9 out of 11 aptamer candidates (Figure S5).

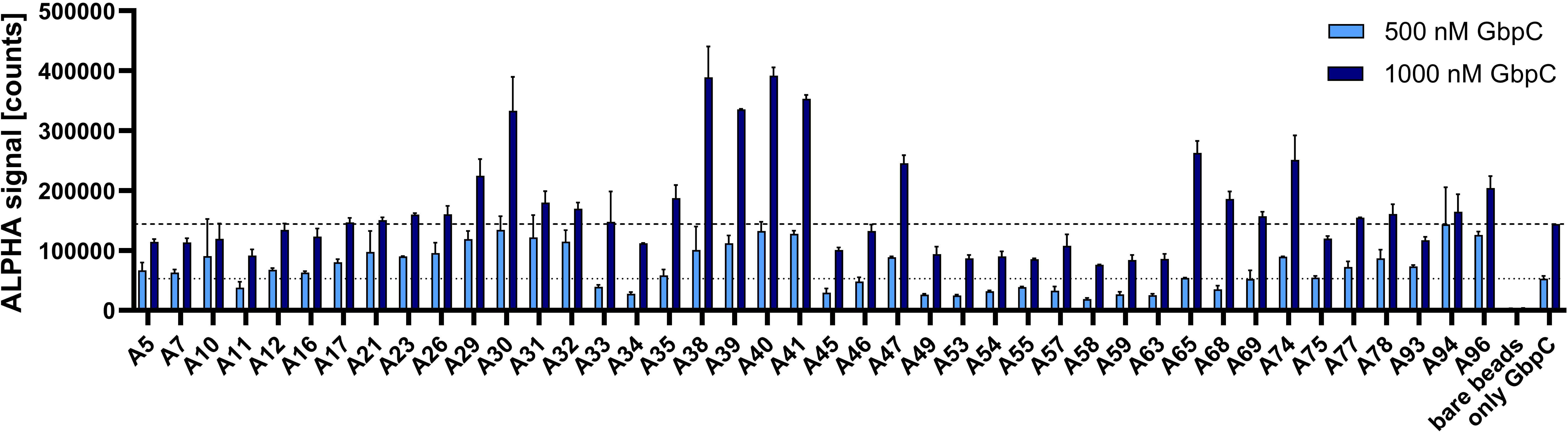

These experiments identified aptamers of the best target protein binding capacity and highlighted the importance of panning the SELEX-derived aptamer candidates.

### 3.3. Binding properties of aptamers

#### 3.2.1. Specificity assay

The best eleven binders were selected for further ALPHA assays to monitor the specificity of each aptamer candidate. In these experiments, the target binding capacity of the aptamers was compared with that of binding to AgI/II, the protein used for counter selection, and AlphaScreen measurements were performed as described in section 2.4.. According to the chemiluminescence signal obtained, all the oligonucleotide sequences studied showed unambiguously higher specificity for the GbpC target molecule, than either structurally analogous AgI/II or bare Ni^2+^-coated acceptor bead controls, and in most cases the measured chemiluminescence was orders of magnitude higher with the target protein, than with AgI/II (Figure 4). These data confirmed the relevance and success of our counter-selection steps using AgI/II and identified the aptamers that were worthy of further study.

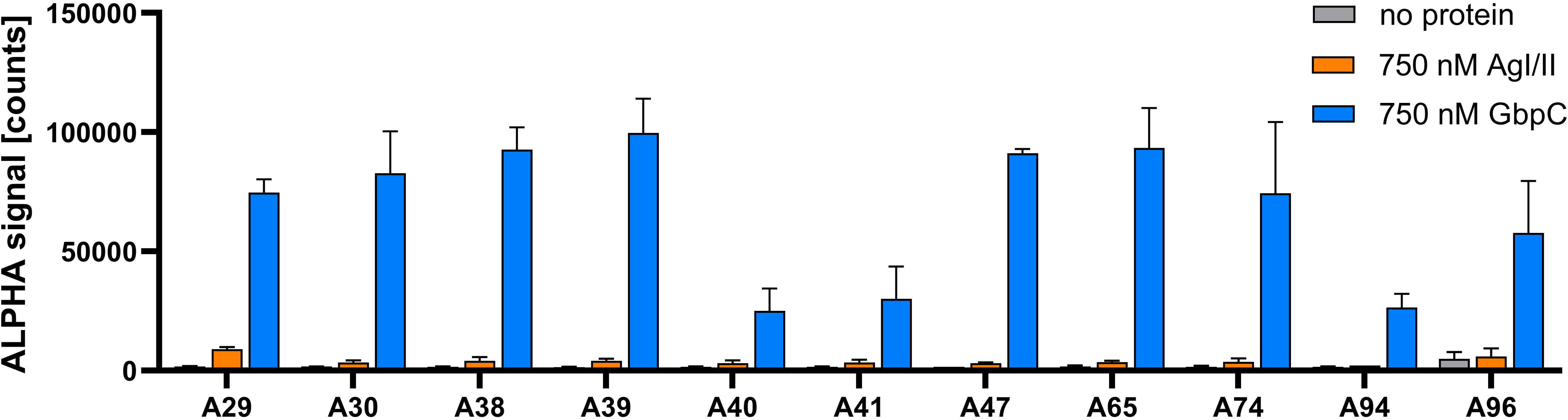

#### 3.2.2. Kinetic analysis

To determine the kinetics of aptamer binding, multiple techniques based on different principles are recommended [59]. We used two distinctly different approaches to measure the kinetics of aptamer-GbpC complex formation. To analyse the dissociation constant (K_D_) of interactions using biolayer interferometry (BLI), one of the interacting partners must be immobilised on the tip surface. To this end, four of the best binding aptamers (A30, A39, A65 and A96) and one of the worst binding aptamers (A58) were chemically synthesised with a 5’-biotin-TEG label and bound to the streptavidin-coated tips. The aptamer-modified sensor tips were immersed in GbpC-containing solutions of different GbpC concentrations, followed by tip washing, and the shift in the interference spectrum was recorded. The data obtained indicated that the rapid association of GbpC with the aptamers is followed by a relatively slow dissociation (Figure S6), resulting in low micromolar dissociation constants in the case of aptamers A30, A39, A65 and A96 (Table 1). The two best performing aptamers, A39 and A96, had the most favourable kinetic parameters with K_D_ values of 5.7 and 0.4 µM, respectively; whereas A58, which was a weak binder according to the AlphaScreen studies, had a dissociation constant more than one order of magnitude higher.

**Table 1.** Characterisation of the kinetics of aptamer-GbpC interaction by biolayer interferometry. The measured association (k_a_) and dissociation (k_d_) rate constants and the calculated equilibrium dissociation constants (K_D_) confirm the interaction between the studied aptamers and GbpC.

| | $k_a$ [1/Ms] | $k_d$ [1/s] | $K_D$ [M] |
| --- | --- | --- | --- |
| **A30** | 1.629e(+3) | 9.974e(-3) | 6.122e(-6) |
| **A39** | 1.472e(+3) | 8.371e(-3) | 5.687e(-6) |
| **A65** | 1.229e(+3) | 9.335e(-3) | 7.598e(-6) |
| **A96** | 5.363e(+3) | 2.362e(-3) | 4.405e(-7) |
| **A58** | 6.47e(+1) | 8.963e(-3) | 1.385e(-4) |

Overall, the BLI analysis showed that most of the aptamers studied formed a stronger complex with the target protein than the GbpC-dextran complex. Furthermore, the BLI data were in agreement with our previous AlphaScreen measurement, indicating the suitability of our aptamer panning approach.

Both of the above methods (ALPHA and BLI) require immobilisation of at least one of the interacting partners; therefore, we decided to provide a further line of evidence for the interaction between the most promising aptamers and GbpC by using microscale thermophoresis (MST). The main advantages of MST are that there is no need for complicated immobilisation protocols as the measurement is performed in any assay buffer or biological fluid, and only a meagre amount of sample is required [60–62]. For the MST studies, the two best performing aptamers (A39 and A96) were synthesised with 5’-Cy5 fluorescent labelling. Both aptamers were produced either with their full sequence or truncated, i.e. with the primer regions removed, in order to determine whether these latter nucleic acid sequences are involved in the formation of the aptamer-protein complex. We mixed all aptamer variants with a dilution series of GbpC in 1xASB + 0.02% Tween-20 and monitored the fluorescence signal.

The determination of the dissociation constants of the protein-aptamer complexes clearly showed that the primer regions of the aptamers play a role in their binding to their target protein, since the dissociation constant of the complexes increased markedly for both aptamers, i.e. from 8.09 to 19.6 µM and from 8.50 to 66.9 µM in the case of A39 and A96, respectively (Figure 5A and 5B). Although the K_D_ value obtained by MST was twenty times higher than that determined by BLI, which seems unexpected, the significant variation in K_D_ values of the same interactions determined by different methods is a documented phenomenon [59,63,64]. Nevertheless, the MST experiments corroborated the GbpC binding capacity of the best performing aptamers and showed that either one or both primer regions of the aptamers are essential components for aptamer-protein complex formation.

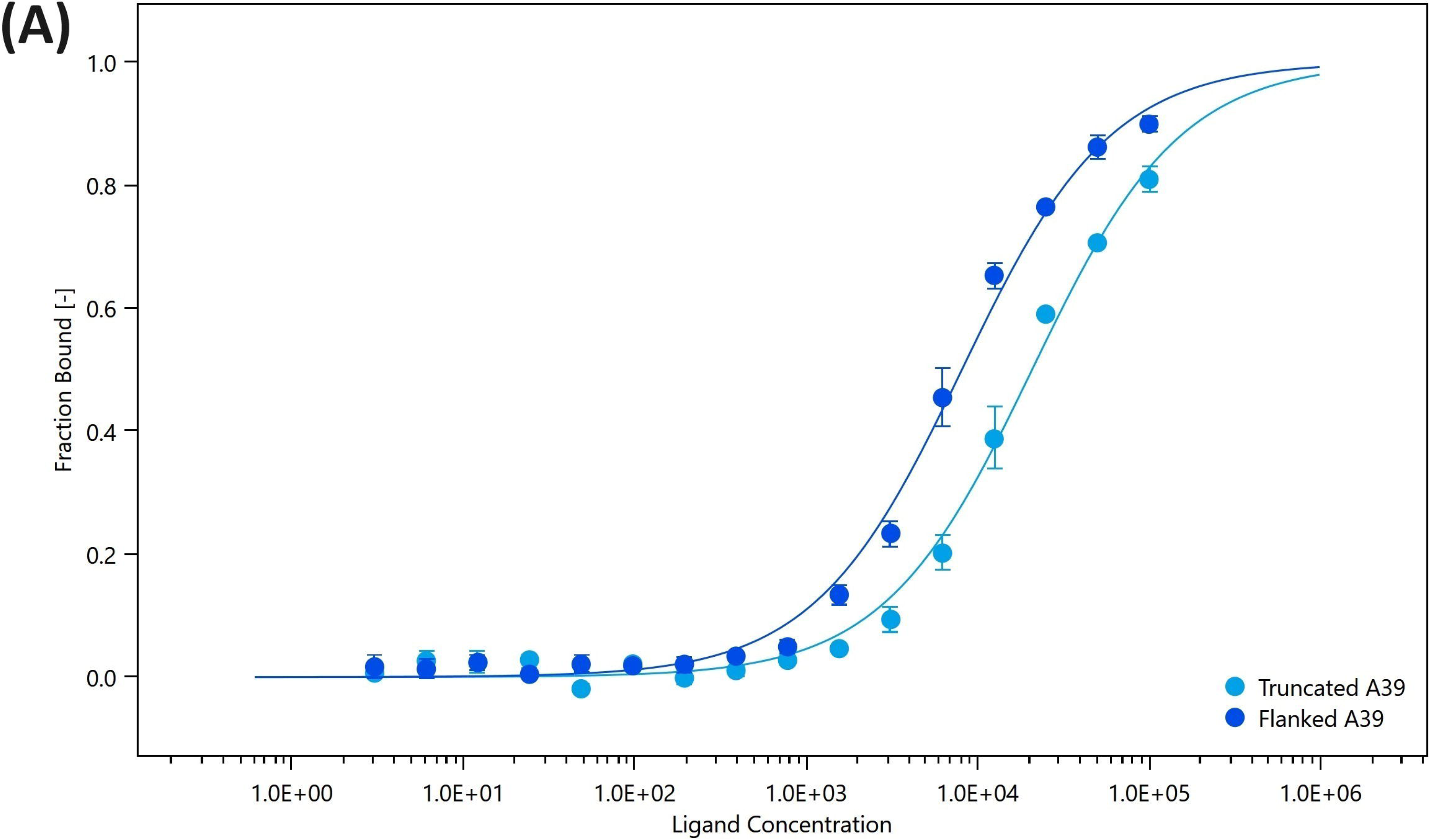
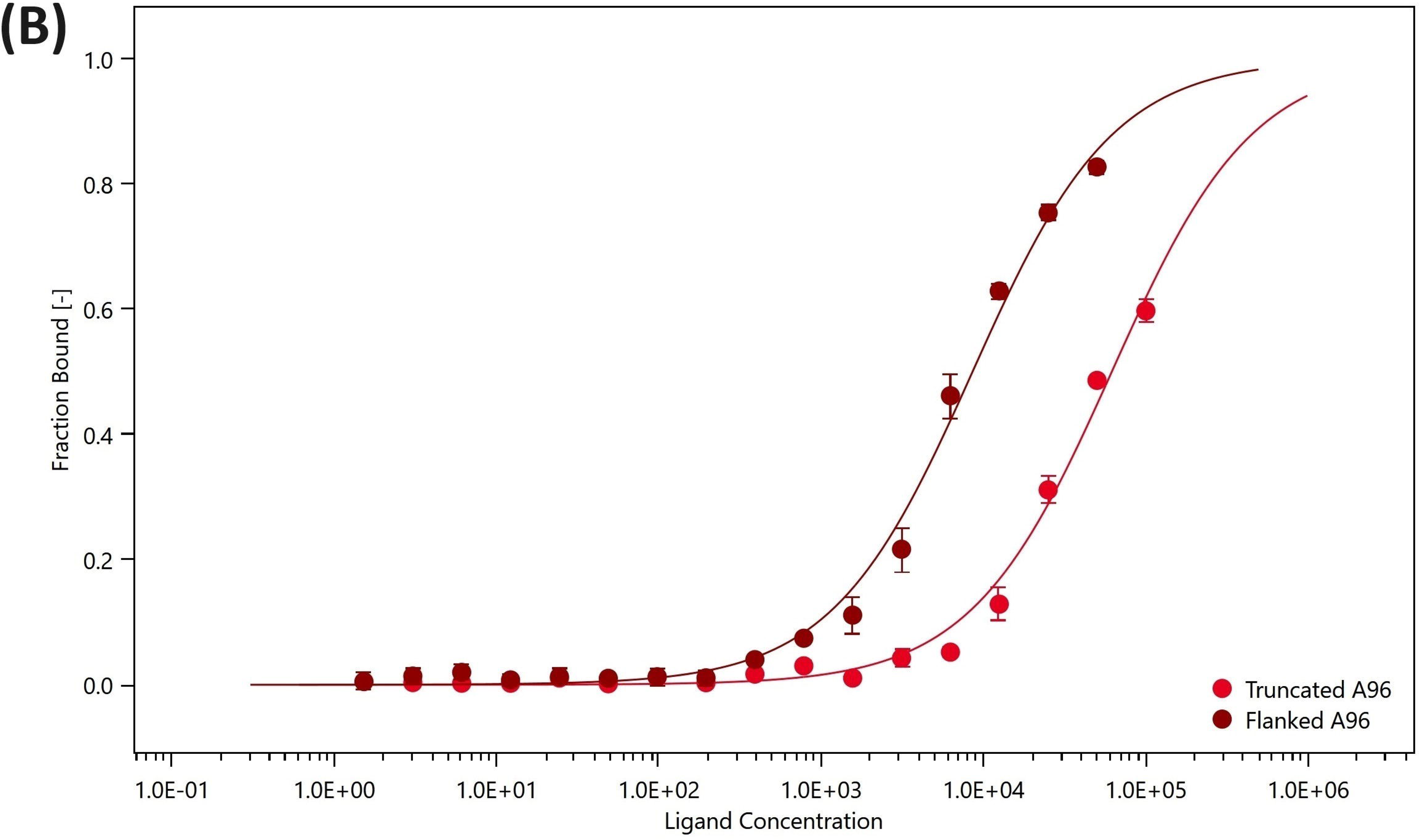
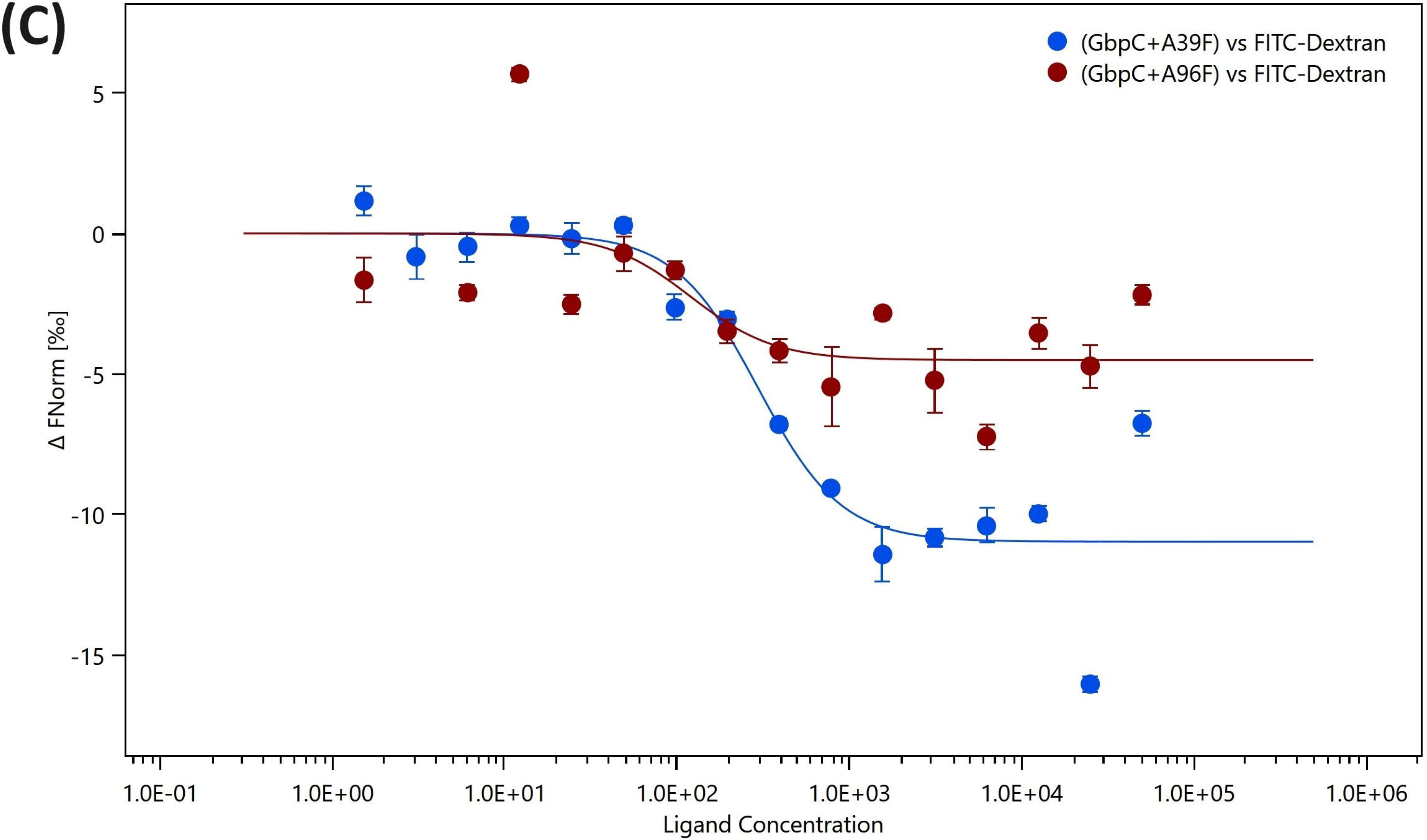

#### 3.2.3. Competitive assay

In our preceding experiments, we demonstrated that both the dextran and the aptamers are capable of binding to GbpC, so we investigated whether they compete for binding to the protein using MST. In competitive binding assays, a constant concentration of Cy5-labelled aptamer and GbpC was pre-incubated and titrated against dextran dilution series. The Hill model was used to fit the F_norm_ curves and determine the half maximal effective concentrations (EC_50_). The EC_50_ value defines the concentration of ligand, in our case dextran, at which 50% of the tracer molecule is bound [65]. The observed slight decrease in the ΔF_norm_ curves and the calculated EC_50_ values of 294 nM and 120 nM for aptamer A39 and A96 respectively (Figure 5C) indicate ligand binding, i.e. both aptamer and dextran compete for GbpC, confirming the inhibitory potential of the tested aptamers. Although these results show that the aptamers can reduce the dextran binding of GbpC, further studies are required to elucidate the mode of inhibition and to identify the exact binding site of the aptamers.

### 3.4. Investigation of biofilm-inhibitory effect of aptamers

After demonstrating the specific binding of GbpC aptamers by various in vitro methods, we investigated the functionality of aptamers in their intended application, i.e. the inhibition of biofilm formation of *S. mutans.* The biofilm inhibitory capacity of aptamers was tested using four good (A30, A39, A65, A96) and one weak (A58) binder. *S. mutans* UA130 and ΔGbpC strains were grown in biofilm-forming medium in the presence of 2 µM of selected unlabelled full-length aptamers. After overnight incubation, biofilm levels were determined by traditional crystal violet staining and the measured absorbance values of the samples were normalised to aptamer-free (untreated) controls.

Aptamers A39 and A96 -the top two performers in the interaction studies - remarkably reduced biofilm formation, when incubated with wild-type *S. mutans* cells. The amount of biofilm was reduced by almost 30% compared to untreated bacterial samples. In contrast, the other two aptamers A30 and A65 with micromolar K_D_ values, similar to the weak binder A58, showed almost no inhibition of biofilm production, when incubated with the wild-type *S. mutans* cell. Importantly, none of these aptamers had a significant inhibitory effect on biofilm formation when the ΔGbpC mutant strain was treated, indicating high specificity of the selected aptamers (Figure 6). These results not only demonstrate the success of aptamer selection, but also highlight the importance of a thorough analysis of the selected aptamers in their intended application to identify the most potent ones.

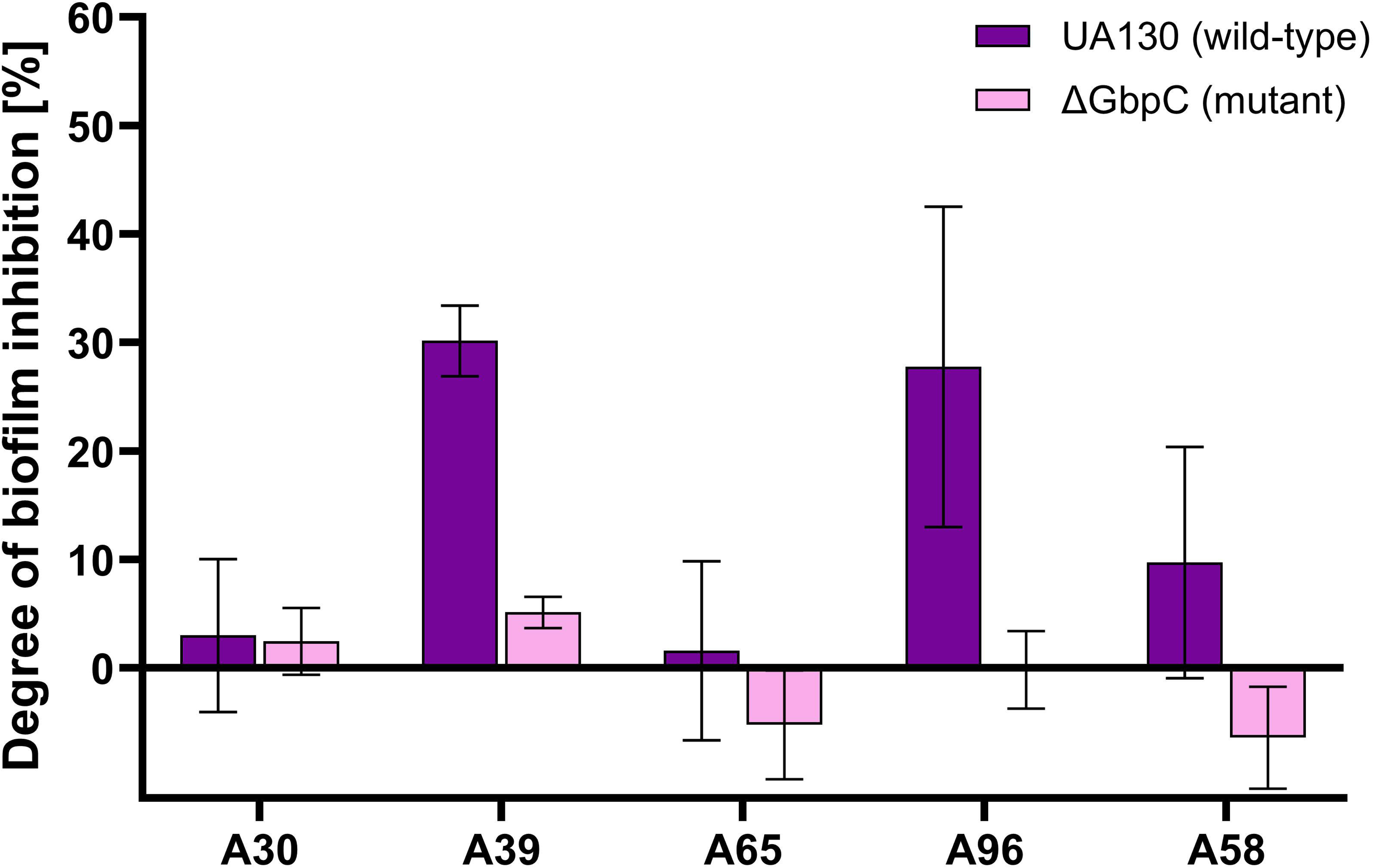

## 4. Conclusion

The aim of our work was to select aptamers against GbpC of *Streptococcus mutans* with biofilm inhibitory potential. The successful production of practical aptamers is basically determined by the use of a suitable target molecule of selection, the implementation of SELEX under the conditions of the intended application of the aptamers and a thorough functional analysis of the candidate aptamers. In order to meet these basic requirements, we demonstrated the proper folding of the extracellular domain of bacterially overexpressed GbpC by analysing its increased spatial stability in the presence of calcium ions and its glucan binding potential with thermal unfolding assay, BLI and MST prior to its application in SELEX. The 3D structure of aptamers is strongly influenced by the prevailing buffer, thus changing the presence and concentration of ions and pH is likely to degrade the aptamer functionality. The long-term intended application of our aptamers is to reduce dental plaque formation. Therefore, the SELEX was implemented in artificial saliva to ensure the correct spatial structure of the aptamers in their practical application.

Although the initial aptamer library is enriched during SELEX, the process can result in tens of thousands of oligonucleotides, and the most enriched oligonucleotides do not necessarily represent the best performing aptamers. We applied a straightforward method, combining AlphaScreen and PBA-PCR, to identify the best target protein binding oligonucleotides. The results illustrated the importance of experimental panning of aptamer candidates, as only about a quarter of the panned oligonucleotides showed clear GbpC binding capacity. Notably, the most abundant oligonucleotide was not identified as a good performer and only one of the oligonucleotides represented in duplicate was among the best performers, further demonstrating the value of experimental panning. Further detailed analysis of the best performing aptamers corroborated applicability of the aptamer panning method used, as all oligonucleotides studied showed selectivity to the target protein over the AgI/II, which was used as a counter selection protein and showed a dissociation constant in the low micromolar range when measured by either BLI or MST.

Finally, we showed that some of the selected aptamers, A39 and A96 had a similar biofilm inhibitory capacity as other previously reported aptamers, i.e. the biofilm formation was reduced by approximately 30% according to crystal violet measurements. Importantly, biofilm formation was not reduced when a GbpC-deficient strain of *S. mutans* was treated with the aptamers. This latter finding provides important evidence for that the selected aptamers are specific for the GbpC and inhibit the biofilm formation by binding to the target protein of SELEX. Given the competitive assay presented, we can hypothesise that the aptamers exert their inhibitory effect by reducing the glucan binding capacity of GbpC, but this assertion requires further experimentation. Somewhat surprisingly, although aptamers A30, A39 and A65 have the same dissociation constant, only A39 was able to inhibit biofilm formation. We suspect that this difference may be due to differences in the stability of the aptamers when exposed to bacterial cells or binding to different domains of GbpC, but these hypothesises have not yet been tested.

The biofilm inhibition potentials of the aptamers presented here are similar to those of the previously reported aptamers, i.e. they fall in the range of 20% to 40% [45,48]. This level of biofilm inhibition may not be sufficient for therapeutics, but the use of aptamers to decorate nanoparticles can significantly increase their efficacy. It has been reported that the functionalisation of graphene oxide with *Salmonella typhimurium* specific aptamers had a synergistic effect and biofilm inhibition was over 90% [66]. A similar effect was described with the combination of *S. mutans* and *P. aeruginosa* selective aptamers with silver nanoparticles and single-walled carbon nanotubes, respectively [67,68]. These results suggest that our aptamers may be of practical importance in the development of dental plaque inhibitors. Moreover, our aptamer selection strategy may be exploited in a range of other medically relevant therapeutic applications in the future.

## 6. CRediT authorship contribution statement

**Ákos Harkai:** Conceptualization, Data curation, Formal analysis, Investigation, Methodology, Project administration, Visualization, Writing – original draft, Writing – review & editing

**Yoon Kee Beck:** Investigation, Validation, Visualization

**Anna Tory:** Investigation, Validation, Visualization

**Tamás Mészáros:** Conceptualization, Funding acquisition, Resources, Supervision, Writing – original draft, Writing – review & editing

## Acknowledgement

We are grateful to Jeffrey A. Banas for providing us kindly the *S. mutans* UA130 and ΔGbpC strains. We express our gratitude to the Institute of Enzymology (HUN-REN) for allowing us to use the MST instrument and to 2bind GmbH (Regensburg) for their expert assistance. We would also like to specially thank Orsolya Dobay for her willing support in the biofilm assays.

## 9. Appendix A: Supplementary data

Please find Appendix A (Supplementary Data) in a separate document.

## 11. Figures and Tables

Please find Figures and their captions in a separate document.

