## Supplementary material for "Selection of streptococcal glucan-binding protein C specific DNA aptamers to inhibit biofilm formation": GbpCapta_captions.docx

**Captions of Figures and Table for**

____________________________________________________________________________________

**Figure 1. Production and functional analysis of GbpC.**

(A) SDS-PAGE shows fractions from the purification steps of GbpC. (M) protein marker, (1) non-induced and (2) auto-induced crude protein extract, (3) IMAC magnetic bead coupled protein, (4) magnetic bead after elution, (5) eluted target protein, (6) buffer-exchanged purified protein.

(B) Analysis of GbpC and biotin-dextran binding by biolayer interferometry. Time-resolved spectral shifts are plotted with increasing protein concentration. The calculated dissociation constant is 13.69 µM.

(C) Analysis of GbpC and FITC-dextran using microscale thermophoresis. The fraction bound is plotted (green) as a function of protein concentration. The calculated dissociation constant is 2.46 µM.

**Figure 2. Schematic of the SELEX procedure used in this study.**

A magnetic bead-immobilised protein fragment of GbpC was used as the target molecule for selection. The non-specifically binding DNA oligonucleotides were removed using AgI/II protein fragment coated magnetic beads and a GbpC protein-deficient *S. mutans* strain. Selection pressure was increased by decreasing the amount of target protein and the incubation time, and by more stringent washing conditions in successive SELEX cycles (created with BioRender.com).

**Figure 3. Panning of candidate aptamers using ALPHAScreen.**

GbpC-coated acceptor beads produce a higher chemiluminescence signal (light and dark blue) with some of the aptamer-coated beads than with the bare streptavidin donor beads, indicating the most promising aptamers (A29, A30, A38, A39, A40, A41, A47, A65, A74, A94, A96). Dashed lines mark the aptamer-free background signals.

**Figure 4. Demonstration of the specificity of the most promising aptamers using AlphaScreen.**

The investigated aptamers clearly produce higher chemiluminescence with GbpC-coated acceptor beads (blue) than with AgI/II-coated (orange) or bare Ni^2+^-coated acceptor beads (grey), indicating that they specifically bind to the target molecule of the selection.

**Figure 5. Analysis of direct and competitive MST data.**

(A) Fraction bound plots of truncated (light blue) and flanked (dark blue) A39 aptamers. The calculated K_D_s of 8.09 and 19.6 µM, respectively.

(B) Fraction bound plots of truncated (light red) and flanked (dark red) A96 aptamers. The calculated K_D_s are 8.50 to 66.9 µM, respectively.

(C) Baseline corrected normalised fluorescence (ΔF_norm_) of competitive assays, resulting EC_50_(A39F)= 294 nM; EC_50_(A96F)= 120 nM.

**Figure 6. Biofilm inhibitory activity of aptamers.**

Crystal violet assays were performed on cells grown in the presence or absence of aptamers to test their biofilm inhibitory activity. Among the sequences tested, aptamers A39 and A96 - the top performers in the interaction studies - reduce biofilm formation in wild-type *S. mutans* (violet), but not in the ΔGbpC strain (rose). The other three aptamers show no inhibitory activity in either strain.

**Table 1. Characterisation of the kinetics of aptamer-GbpC interaction by biolayer interferometry.**

The measured association (k_a_) and dissociation (k_d_) rate constants and the calculated equilibrium dissociation constants (K_D_) confirm the interaction between the studied aptamers and GbpC.
