## Supplementary material for "Selection of streptococcal glucan-binding protein C specific DNA aptamers to inhibit biofilm formation": GbpCapta_supplementary.docx

____________________________________________________________________________________

**This document includes:**

Supplementary Table 1.	Components of a commercial Artificial Saliva without mucin.

Supplementary Table 2.	Detailed protocol for applied SELEX procedure.

Supplementary Table 3.	List of applied oligonucleotides.

Supplementary Method 1. Thermal Unfolding Assay of GbpC

Supplementary Figure 1.	Thermal unfolding assay of GbpC in different buffers.

Supplementary Figure 2.	Primary structure alignment of GbpC^111-522^ and AgI/II^457-993^	protein fragments.

Supplementary Figure 3.	Illustration of variable regions of aptamer sequences and their	prevalence.

Supplementary Figure 4.	Glucan-binding function of proteins with ALPHA donor bead.

Supplementary Figure 5.	In-silico analysis of the best performing aptamer candidates of	AlphaScreen measurement.

Supplementary Figure 6.	Binding curves of A30, A39, A65, A96, A58 aptamers measured by	biolayer interferometry.

**Supplementary Table 1. Components of a commercial Artificial Saliva without mucin.**

| **Component** | **Concentration** | | **Molecular weight** |
| --- | --- | --- | --- |
| Sodium Chloride | 0.88 g/L | 15.06 mmol/L | 58.44 g/mol |
| 0.2 M Potassium Phosphate Monobasic | 7.7 mL/L | 1.54 mmol/L | 136.1 g/mol |
| 0.2 M Potassium Phosphate Dibasic | 12.3 mL/L | 2.46 mmol/L | 174.2 g/mol |
| Potassium Chloride | 1.04 g/L | 13.95 mmol/L | 74.55 g/mol |
| Potassium Thiocyanate | 0.19 g/L | 1.96 mmol/L | 97.2 g/mol |
| Calcium Chloride monohydrate | 0.13 g/L | 1.01 mmol/L | 129 g/mol |
| Magnesium Chloride heptahydrate | 0.04 g/L | 0.18 mmol/L | 221.2 g/mol |
| Ammonium Chloride | 0.11 g/L | 2.06 mmol/L | 53.5 g/mol |
| Sodium Bicarbonate | 0.42 g/L | 5.00 mmol/L | 84 g/mol |
| Urea | 0.12 g/L | 2.00 mmol/L | 60,1 g/mol |
| ProClin™ 300 (preservative) | 0.3 mL/L | 1.17 mmol/L | 264.8 g/mol |
| pH = | 7.00 | |  |

**Supplementary Table 2. Detailed protocol for applied SELEX procedure.**

| **Round** | **Target amount** | **Binding buffer** | **Incubation time and temperature** | **Washing buffer** | **Washing time** |
| --- | --- | --- | --- | --- | --- |
| **1** | 600 pmol GbpC | 1xASB | 60 min, 25°C | 1x 300 µl 1xASB | 1x 5 min |
| **2** | 450 pmol GbpC | 1xASB  +0.01 µg/ml salmon sperm DNA | 60 min, 25°C | 2x 300 µl 1xASB | 2x 5 min |
| **3** | 300 pmol GbpC | 1xASB  +0.01 µg/ml salmon sperm DNA | 30 min, 25°C | 2x 300 µl 1xASB,  1x 300 µl 1xASB  +0.01% Tween20 | 3x 5 min |
| **C1** | 600 pmol AgI/II | 1xASB | 30 min, 25°C | - | - |
| **4** | 600 pmol GbpC | 1xASB  +0.1 µg/ml salmon sperm DNA | 30 min, 25°C | 3x 300 µl 1xASB | 3x 5 min |
| **5** | 300 pmol GbpC | 1xASB  +0.1 mg/ml mucin  +0.1 µg/ml salmon sperm DNA | 30 min, 25°C | 3x 300 µl 1xASB  +0.01% Tween20 | 3x 10 min |
| **C2** | approx. 8x10^8^ CFU/ml  *S. mutans* ΔGbpC cells | 1xASB | 15 min, 25°C | - | - |
| **6** | 600 pmol GbpC | 1xASB  +0.1 µg/ml salmon sperm DNA | 30 min, 25°C | 3x 300 µl 1xASB  +0.05% Tween20 | 3x 5 min |
| **7** | 300 pmol GbpC | 1xASB  +0.1 mg/ml mucin  +0.1 µg/ml salmon sperm DNA | 30 min, 25°C | 3x 300 µl 1xASB  +0.1 mM dextran sulfate | 3x 5 min |
| **C3** | 600 pmol AgI/II | 1xASB | 30 min, 25°C | - | - |
| **8** | 300 pmol GbpC | 1xASB  +0.1 mg/ml mucin  +0.5 µg/ml salmon sperm DNA | 45 min, 25°C | 3x 300 µl 1xASB  +0.1 mM dextran sulfate | 3x 10 min |

**Supplementary Table 3. List of applied oligonucleotides.**

| **Name** | **Sequence (5'**🡪**3')** | **Manufacturer** |
| --- | --- | --- |
| Cloning forward primer (M13F) | GTAAAACGACGGCCAG | Thermo |
| Cloning reverse primer (M13R) | CAGGAAACAGCTATGAC | Thermo |
| cPCR forward primer (193) | AAACGACGGCCAGTGAATTG | Eurofins |
| cPCR reverse primer (194) | TTATGCTTCCGGCTCGTATG | Eurofins |
| DNA Aptamer Library (Lib13) | AGATACCAATACGCTGCC  -(N40)-  GCCACTGGTAACGACATC | Sigma |
| Lib13 forward primer | AGATACCAATACGCTGCC | Eurofins |
| Lib13 reverse primer | GATGTCGTTACCAGTGGC | Eurofins |
| Lib13 reverse complement | GCCACTGGTAACGACATC | Eurofins |
| **Synthesized aptamers** | | |
| A30 | AGATACCAATACGCTGCC-TTCTTGAAAAAGGGTAGCGCAGGCCGCTAATGACATAACT-GCCACTGGTAACGACATC | Eurofins |
| A39 | AGATACCAATACGCTGCC-TAGCCCCTACCCTGTTGCTCCCCCTCGCCCCTCTCACCCC-GCCACTGGTAACGACATC | Eurofins |
| A65 | AGATACCAATACGCTGCC-TTTCCAATTACGCCATGCATCACCCCTAAAATCTGAATAC-GCCACTGGTAACGACATC | Eurofins |
| A96 | AGATACCAATACGCTGCC-CCGCCGGTCCCGAGCTACAACACTTTACGCCCCCCTCCCC-GCCACTGGTAACGACATC | Eurofins |
| A58 | AGATACCAATACGCTGCC-ATACCCACCCATCACCGAGTCGACACTCACCTAACAGCCG-GCCACTGGTAACGACATC | Eurofins |
| A39 truncated | TAGCCCCTACCCTGTTGCTCCCCCTCGCCCCTCTCACCCC | Eurofins |
| A96 truncated | CCGCCGGTCCCGAGCTACAACACTTTACGCCCCCCTCCC | Eurofins |

**Supplementary Method 1. Thermal Unfolding Assay of GbpC**

Brief protocol for Thermal Unfolding Assay of GbpC

The thermal stability of spatial structure of purified GbpC was investigated using GloMelt^TM^ Thermal Shift Protein Stability Kit (Biotium) according to manufacturer's protocol. 10 µl of reaction mixture comprised of 1x GloMelt dye, 50 nM ROX reference dye and 10 µg GbpC diluted with one of the test buffers (PBS, PBS +1 mM CaCl_2_, Artificial Saliva Buffer without mucin). Triplicates were dispensed in MicroAmp^Ⓡ^ Fast 96-Well Reaction Plate (Applied Biosystem) and protein unfolding was monitored by QuantStudio 12K Flex Real-Time PCR System (Applied Biosystem).

Distinct peak appears on the melting curve of GbpC in the presence of calcium-ion containing test buffers, indicating the formation of a more stable domain in the tertiary structure. This observation supports that the overexpressed protein retains its functional structure even after purification.

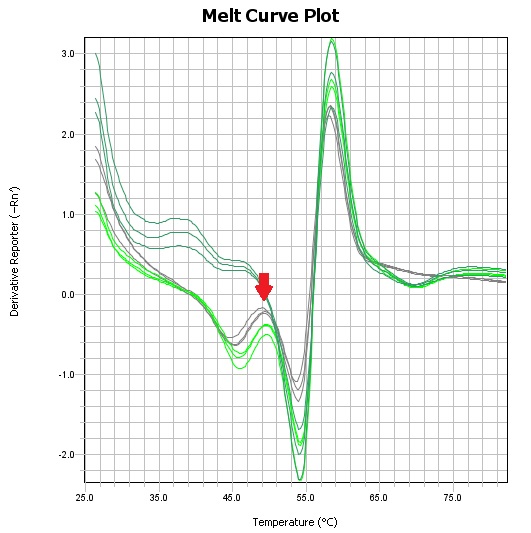

**Supplementary Figure 1. Thermal unfolding assay of GbpC in different buffers.** A distinct peak (red arrow) appears at 49°C in calcium supplemented PBS (light green) and calcium containing artificial saliva buffer (grey), but not in PBS (dark green), indicating increased spatial structural stability of GbpC in the presence of calcium.

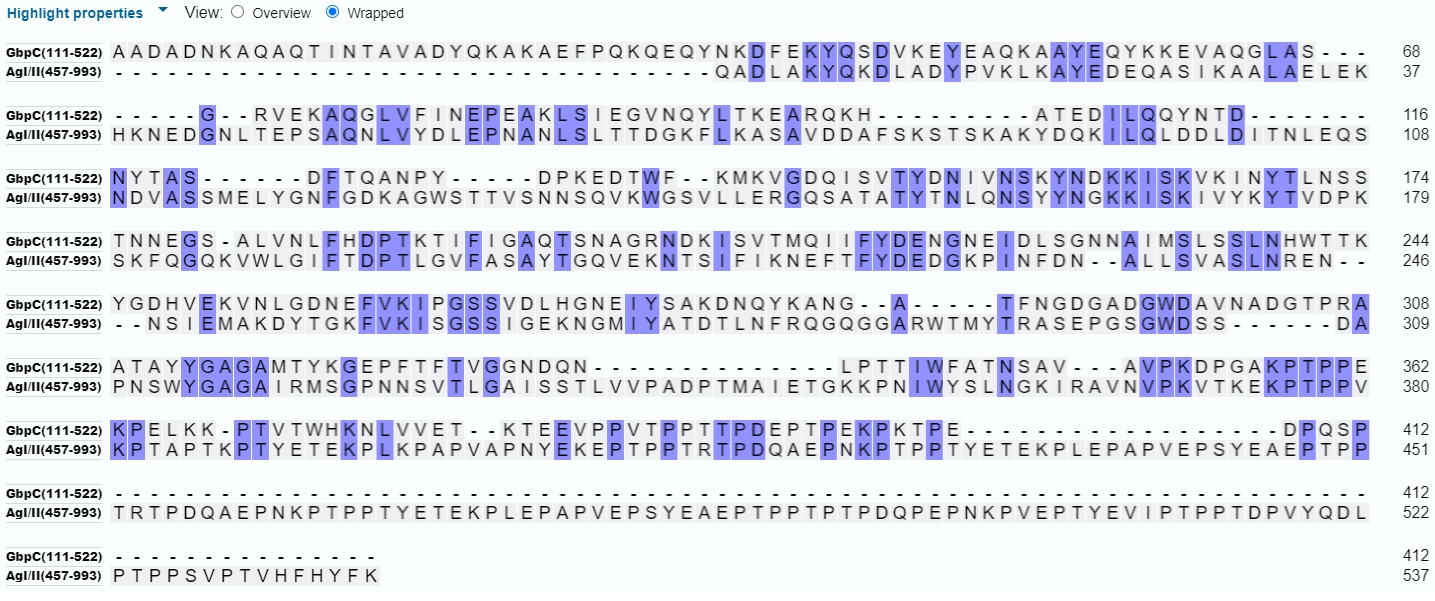

**Supplementary Figure 2. Primary structure alignment of GbpC^111-522^ and AgI/II^457-993^ protein fragments.** Identical amino acids are highlighted in lilac.

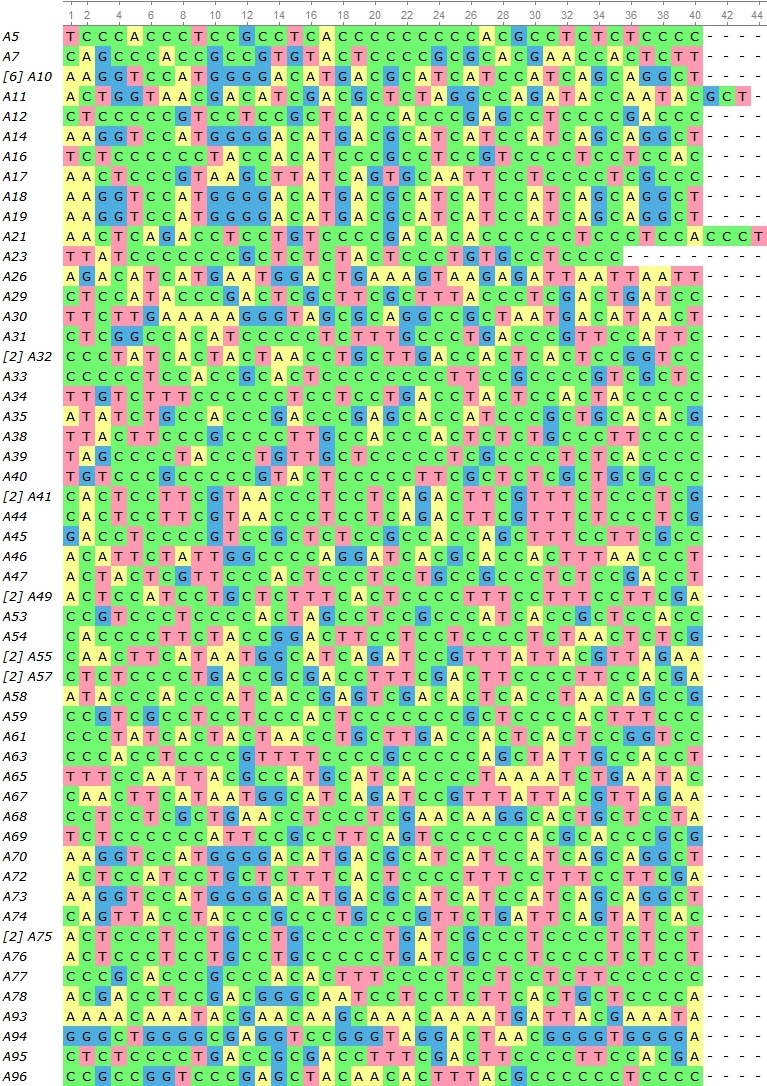

**Supplementary Figure 3. Illustration of variable regions of aptamer sequences and their prevalence (indicated in square brackets in front of the sequence name).**

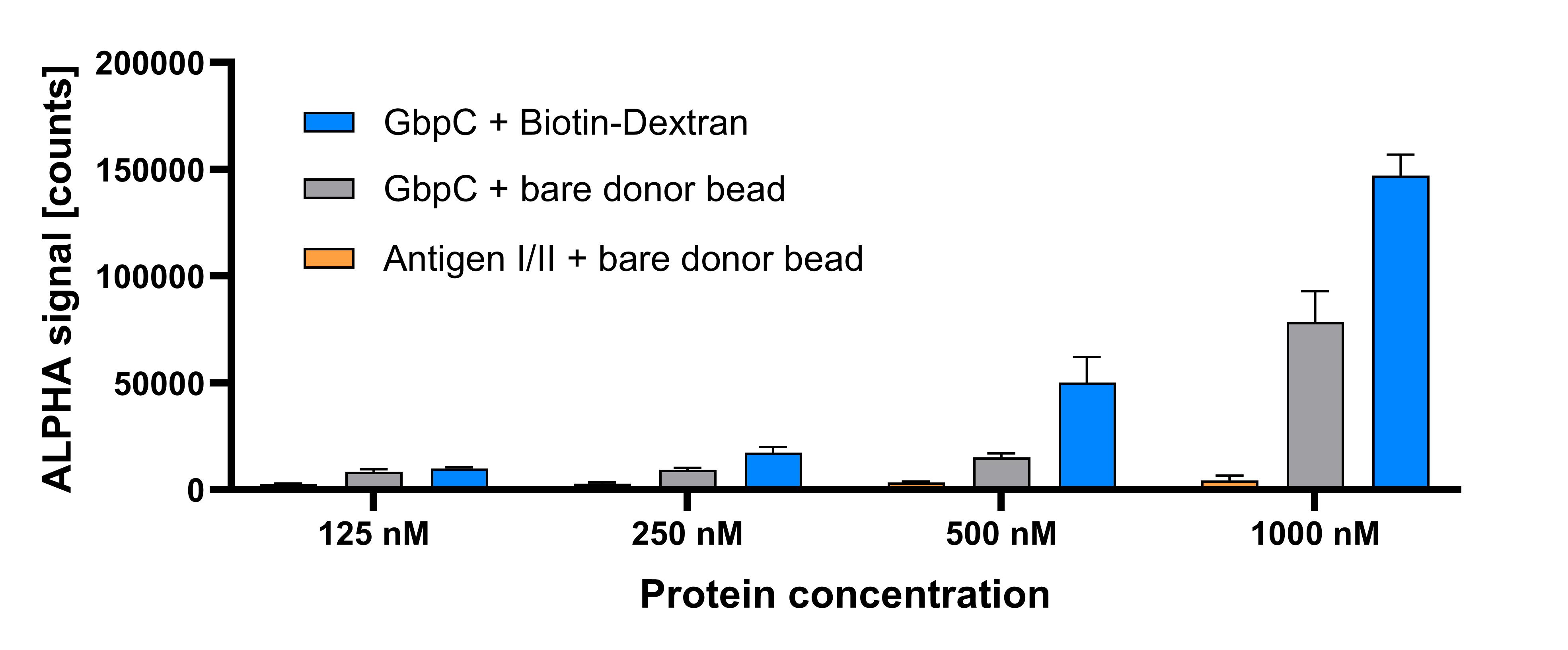
**Supplementary Figure 4. Glucan-binding function of proteins with ALPHA donor bead.**

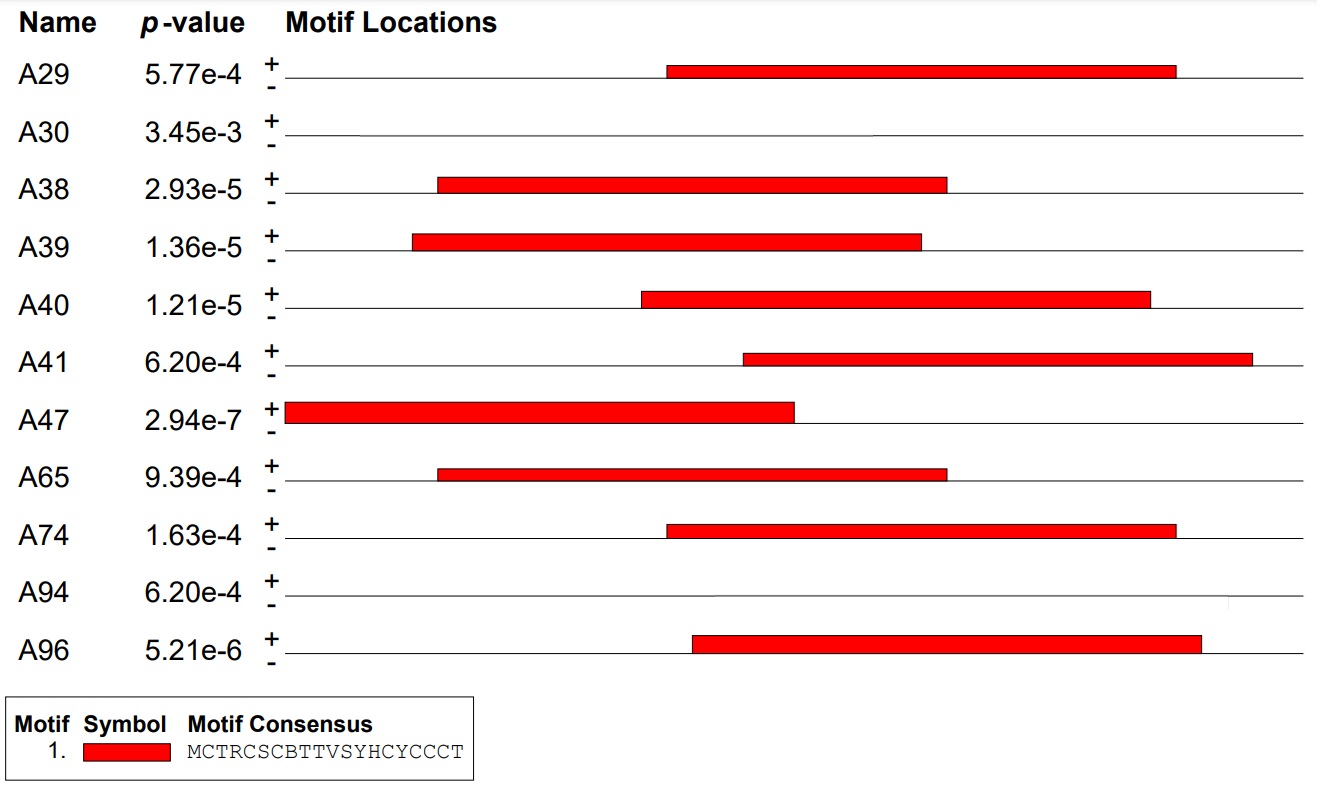

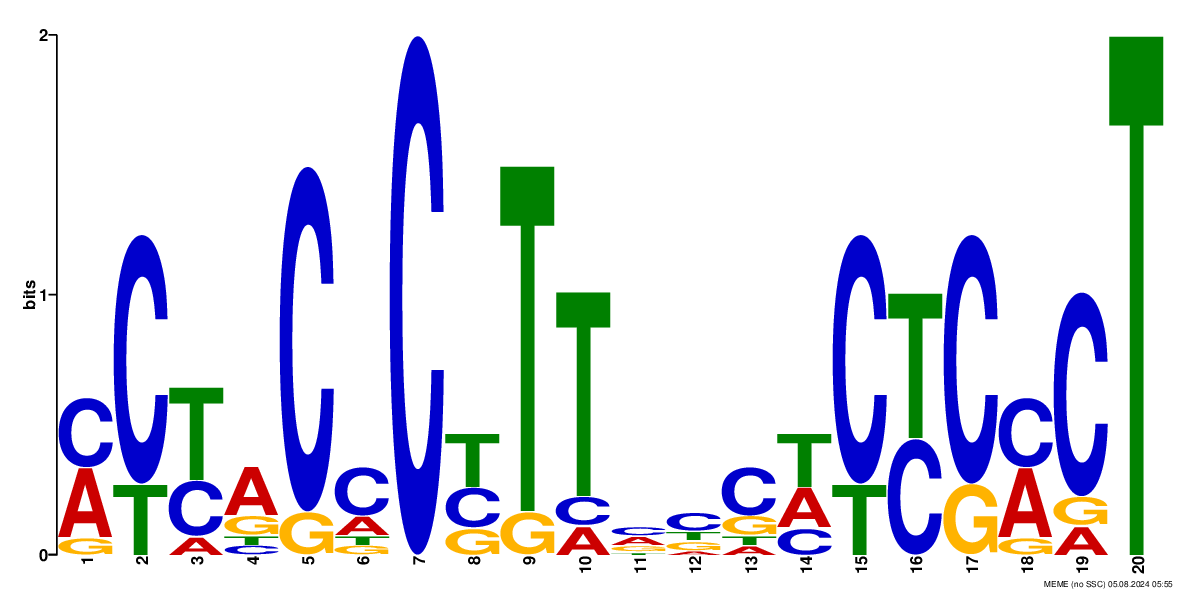

**Supplementary Figure 5. In-silico analysis of the best performing aptamer candidates of AlphaScreen measurement.** The most frequent C-rich motif (red line) was found in almost all sequences using online motif-based sequence analysis tool (MEME). Nucleic acid sequence of the motif is given in the box.

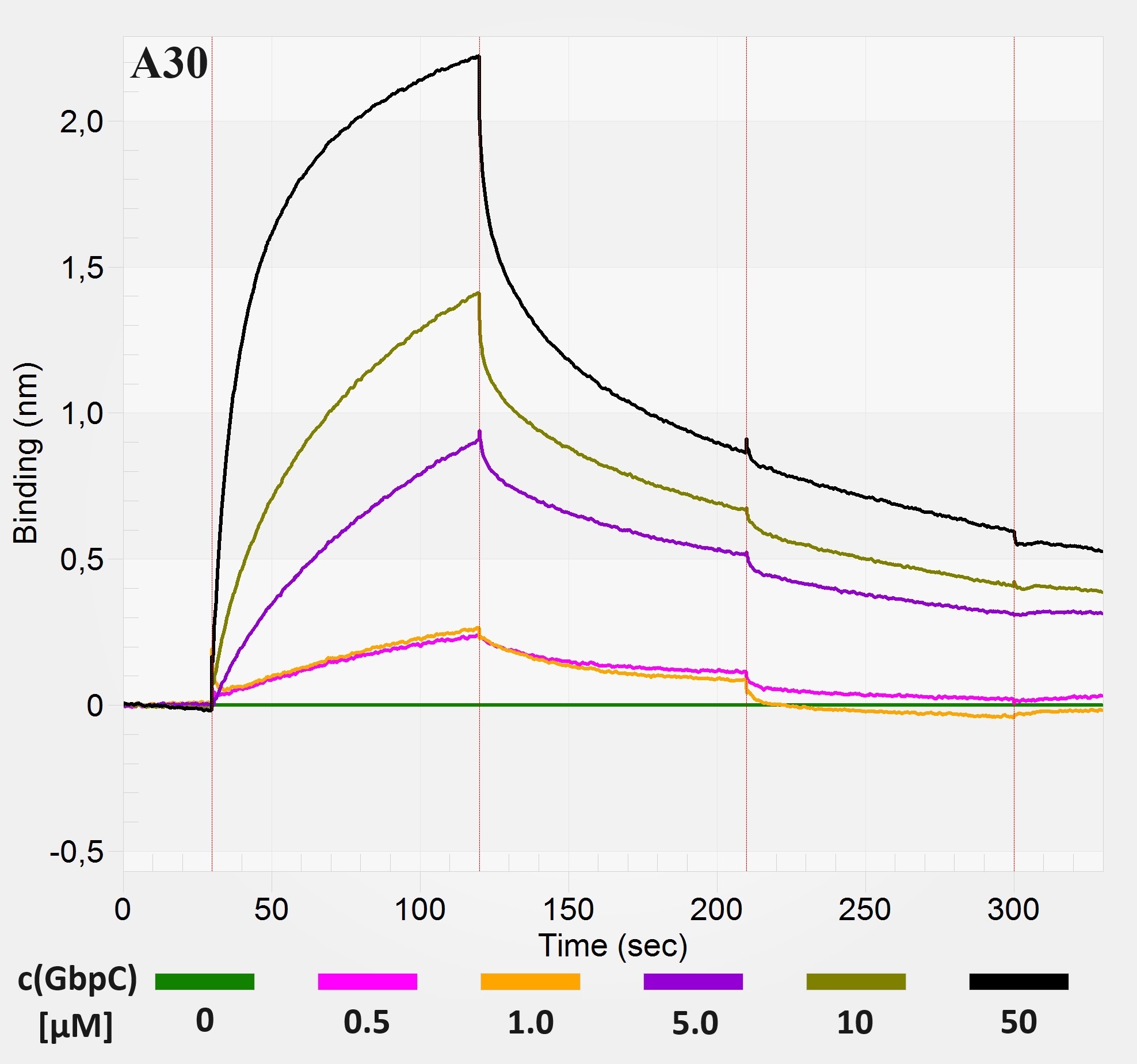

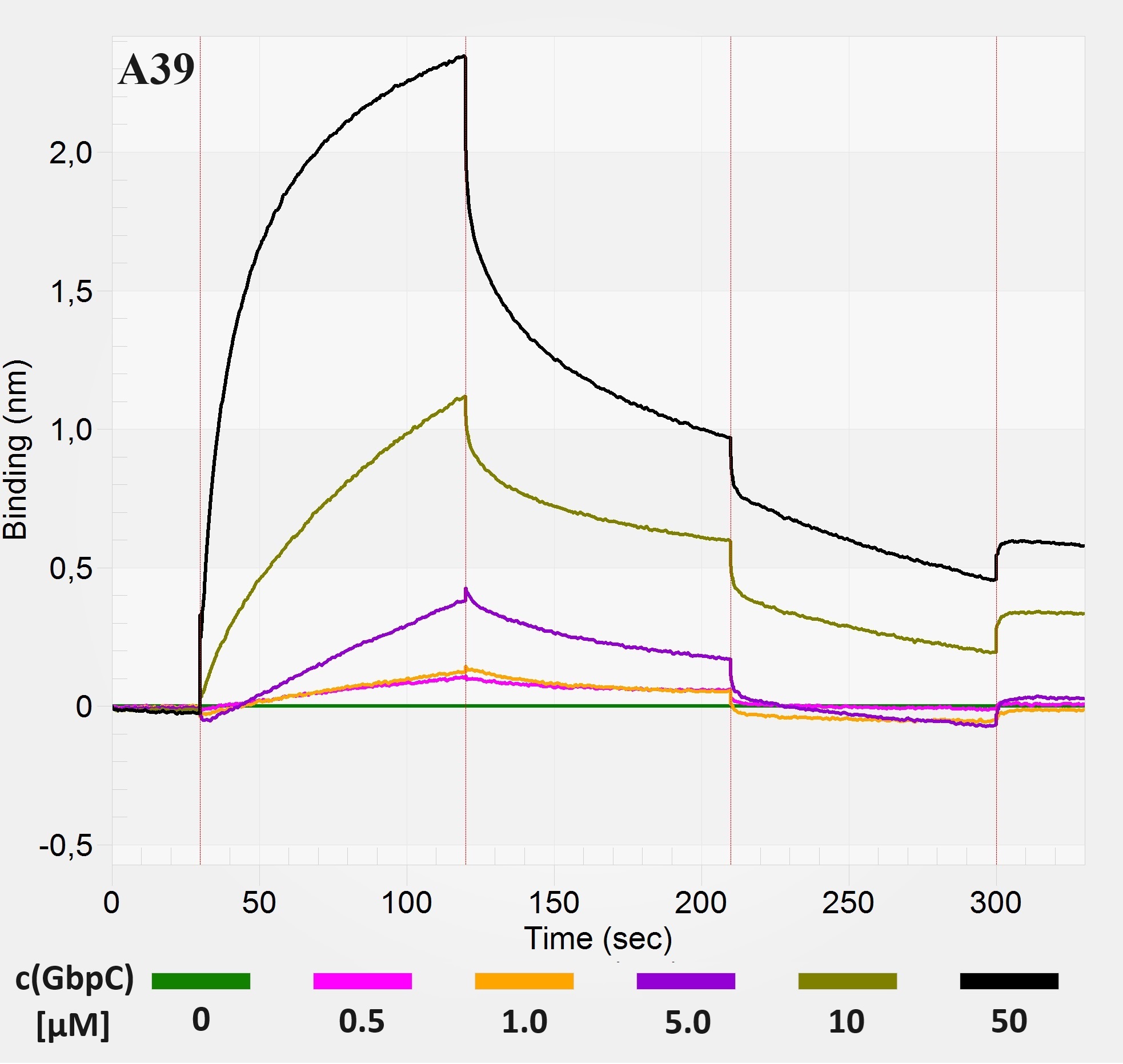

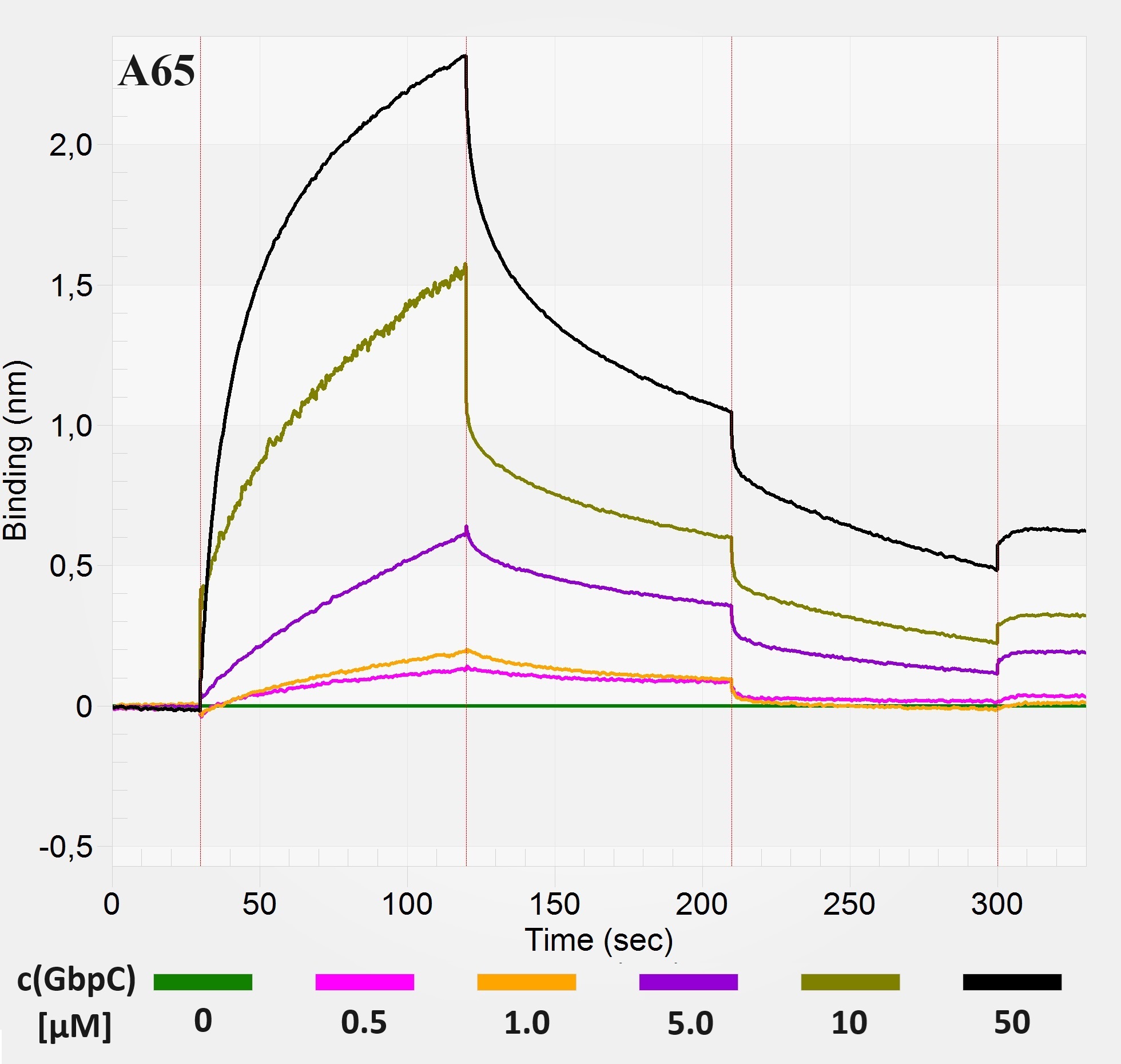

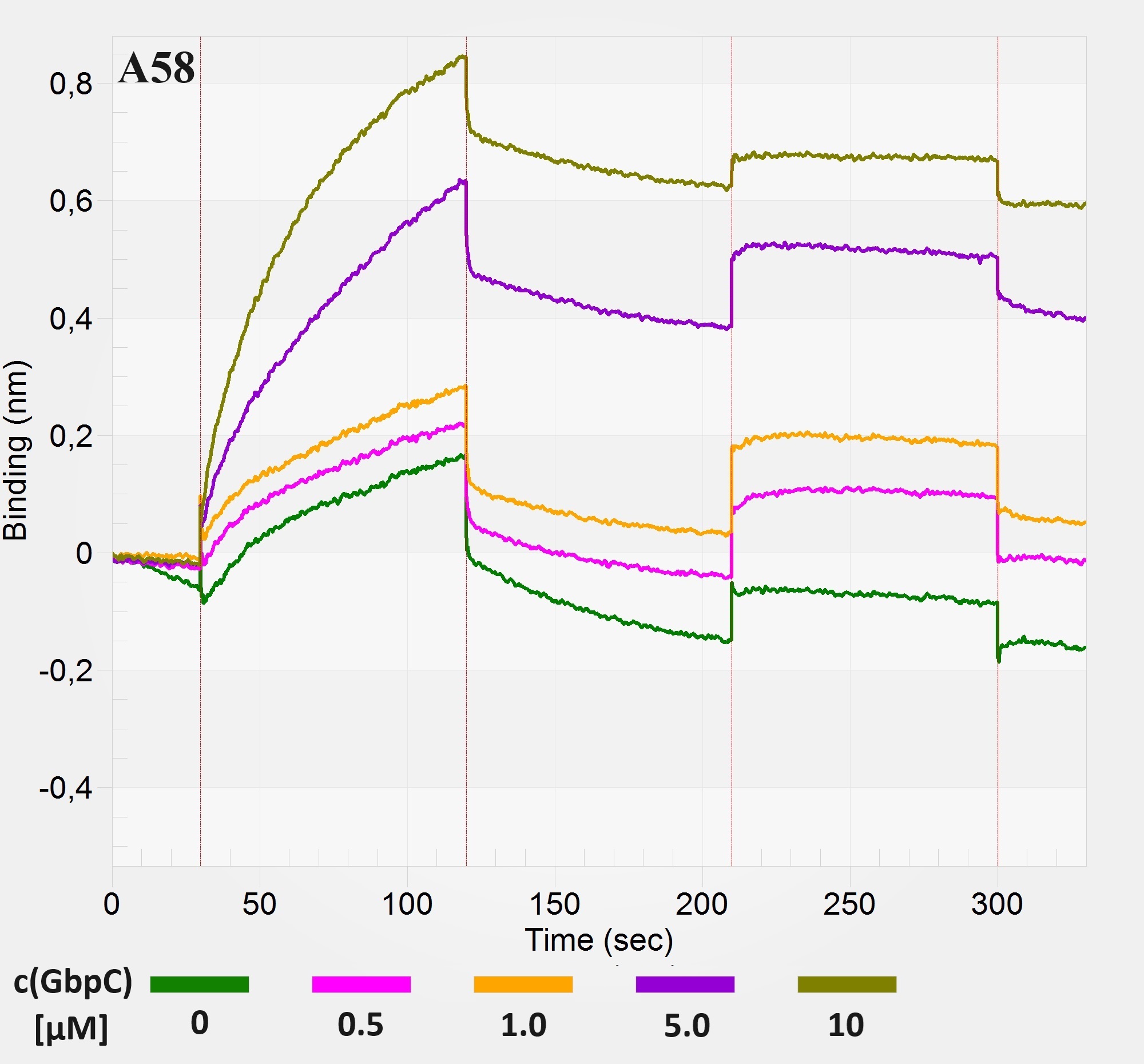

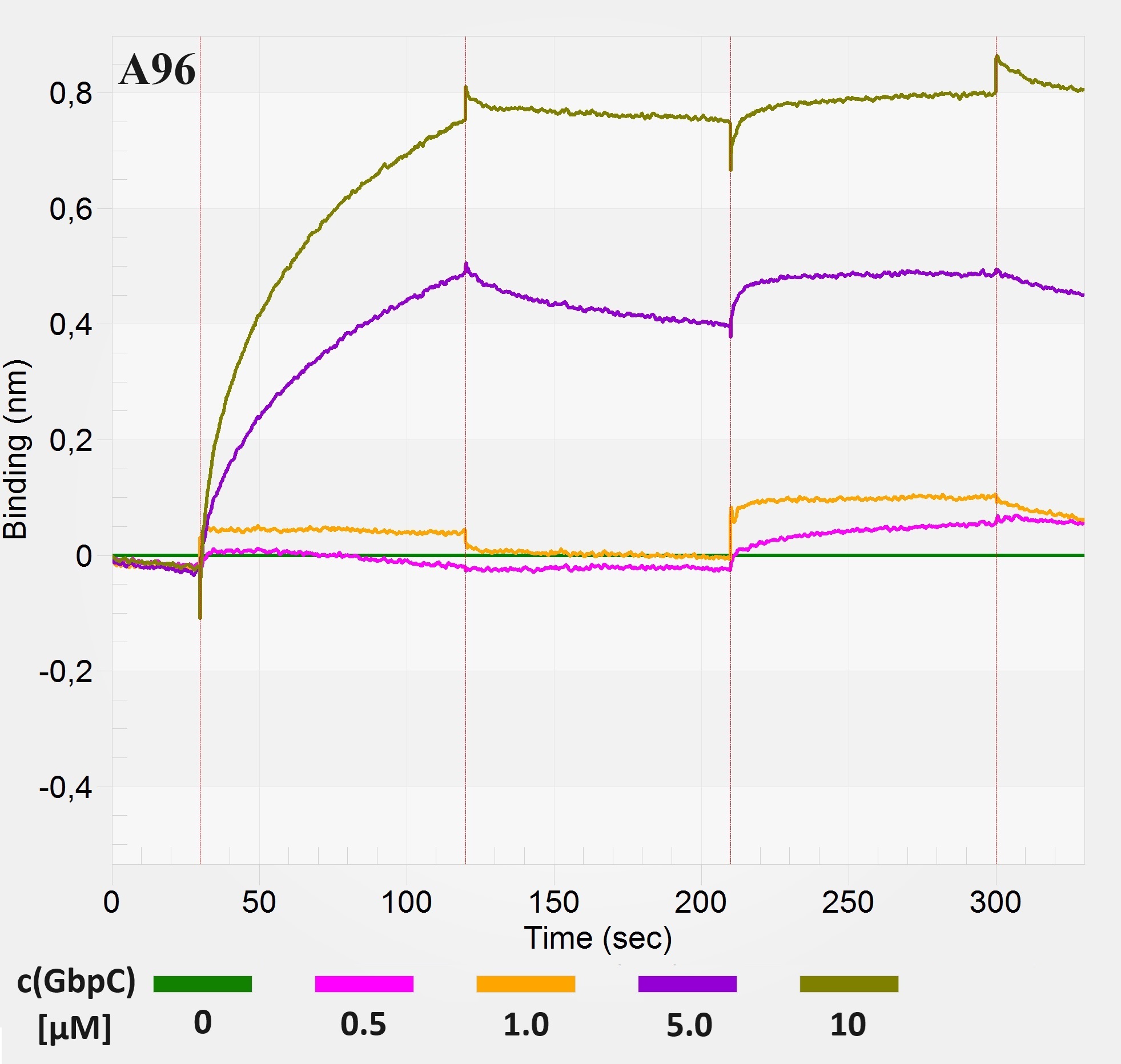

**Supplementary Figure 6. Binding curves of A30, A39, A65, A58, A96 aptamers measured by biolayer interferometry.** Following loading of the streptavidin-coated sensor, the initial baseline (30 s), association (90 s), dissociation (90 s), wash (90 s) and final baseline (30 s) steps were measured.
